# Conserved Neuronal-like and Secretory Programs Define the Spatial Architecture of Gastroenteropancreatic Neuroendocrine Tumors

**DOI:** 10.64898/2025.12.28.696762

**Authors:** Julie Karam, Samantha E. Hoffman, Amanda Garza, Dan Gui, Hannah I. Hoffman, Breanna M. Titchen, Yutaro Tanaka, Erica Pimenta, Theodora Pappa, Laura Valderrabano, Kevin Bi, Riaz Gillani, Lauren Brais, Erin Shannon, Jason L. Hornick, Jihye Park, Jennifer Chan, Eliezer M. Van Allen

## Abstract

Gastroenteropancreatic neuroendocrine tumors (GEP-NETs) are clinically heterogeneous malignancies whose biology and microenvironmental organization remain poorly understood. Here, we integrated single-nucleus multiomic (snRNA-seq and snATAC-seq) and spatial transcriptomic profiling across 38 well-differentiated pancreatic (pNET) and small-intestinal (siNET) tumors to define conserved malignant programs, their regulatory circuits, and spatial niches. We observed two conserved malignant cell programs spanning a continuous transcriptional spectrum: a neuronal-like program (si-cNMF1/p-cNMF1), and a secretory neuroendocrine program (si-cNMF2/p-cNMF2). Matched chromatin accessibility profiles uncovered distinct, tissue-specific regulatory networks, including *MAX::MYC* and *MITF* transcription factor binding motifs in siNETs versus *ISL1* and *TFAP4* in pNETs, indicating organ-specific epigenetic control. Spatial transcriptomic analyses revealed that si/p-cNMF1-high regions localized to high cell density, immune-rich tumor areas, whereas si/p-cNMF2-high regions occupied stromal and vascularized niches and co-occured with fibroblast and endothelial compartments enriched for *TGFB1-ITGB1*, *VEGFA-FLT1*, and *LAMA2-ITGA1* signaling. Across both tumor types, the cNMF2 program was enriched in metastatic lesions and was enrichedfor pro-fibrotic and pro-angiogenic gene signatures. Thus, GEP-NETs are organized along a conserved neuronal-to-secretory axis defined by distinct epigenetic programs and spatially coupled to specific microenvironmental niches. This framework unifies NET heterogeneity across organ sites and identifies pathway-specific, microenvironment-linked vulnerabilities for therapeutic targeting.

## INTRODUCTION

Well-differentiated neuroendocrine tumors (NETs) are rare malignancies that arise from secretory cells of the neuroendocrine system and can affect multiple organ sites. Gastroenteropancreatic NETs (GEP-NETs), which originate in the gastrointestinal tract or pancreas, account for nearly two-thirds of all NET diagnoses (1). Their incidence has risen nearly five-fold in the United States over the past several decades, a trend attributed in part to advances in imaging and endoscopy but also reflecting the increasing prevalence of disease in an aging population (2). As a result, solving the clinical challenges posed by GEP-NETs is therefore an increasingly urgent priority.

Despite the increasing incidence, therapeutic advances for GEP-NETs have lagged. Surgical resection remains the only curative option for localized disease, whereas medical therapies typically slow progression without inducing durable remission (3). A major barrier to therapeutic development is the incomplete understanding of GEP-NET biology compared to other gastrointestinal cancers (4). Their indolent growth has hampered the generation of representative preclinical models, and their rarity has limited the size of clinical cohorts and available samples for characterization. Direct high-resolution profiling of primary human samples therefore represents a powerful strategy to overcome these limitations and reveal new insights into GEP-NET pathophysiology.

Recent single-cell transcriptomic studies of GEP-NETs and their microenvironment have identified immunosuppressive myeloid populations with therapeutic potential or tumor-intrinsic malignant programs (5–7), though the small cohort sizes,heterogeneous sequencing approaches, and incomplete assessment of the microenvironment of these studies have limited generalizability. The spatial architecture of GEP-NETs, particularly their prominent fibrosis and vascular remodeling that may indicate novel biological underpinnings and therapeutic opportunities, has also not been comprehensively examined (8). At the same time, although GEP-NETs appear morphologically homogeneous, their diverse clinical behaviors suggest malignant cell transcriptional or epigenetic heterogeneity that may underlie subtleties in histology and have clinical relevance. Here, we hypothesized that integrating single-cell multiomics with spatial transcriptomic profiling of pancreatic and small intestinal NETs could elucidate previously unappreciated heterogeneity of malignant cell states and define their interactions with the surrounding stroma and immune microenvironment. Single-cell and spatial technologies now allow dissection of tumor ecosystems at unprecedented resolution, and their application to NETs offers the opportunity to uncover conserved transcriptional programs, epigenetic regulatory networks, and spatial niches that underlie progression and metastasis.

To this end, we present the largest multi-modal study of GEP-NETs to date, combining single-nucleus, and spatial transcriptomic approaches to define conserved transcriptional programs across samples and tumor types, and to uncover their epigenetic regulators, spatial organization, and microenvironmental interactions that may have biological and clinical significance.

## RESULTS

### Multi-omic analysis of pancreatic and small intestinal NETs at single-nuclei resolution

We performed joint single-nuclei RNA-sequencing (snRNA-seq) and assay for transposase-accessible chromatin (snATAC)-seq on 38 archival flash-frozen tumor samples of well-differentiated NETs originally resected at Brigham and Women’s Hospital between 2015 and 2021, consisting of 21 pancreatic NETs (pNETs) and 17 small intestinal NETs (siNETs) (Figure 1A, Methods). In addition, high-quality formalin-fixed, paraffin-embedded (FFPE) slides from pNET and siNET samples were used to generate spatial transcriptomic sequencing data (n = 19) using the 10x Visium platform (Figure 1A, Methods).

**Figure 1:**
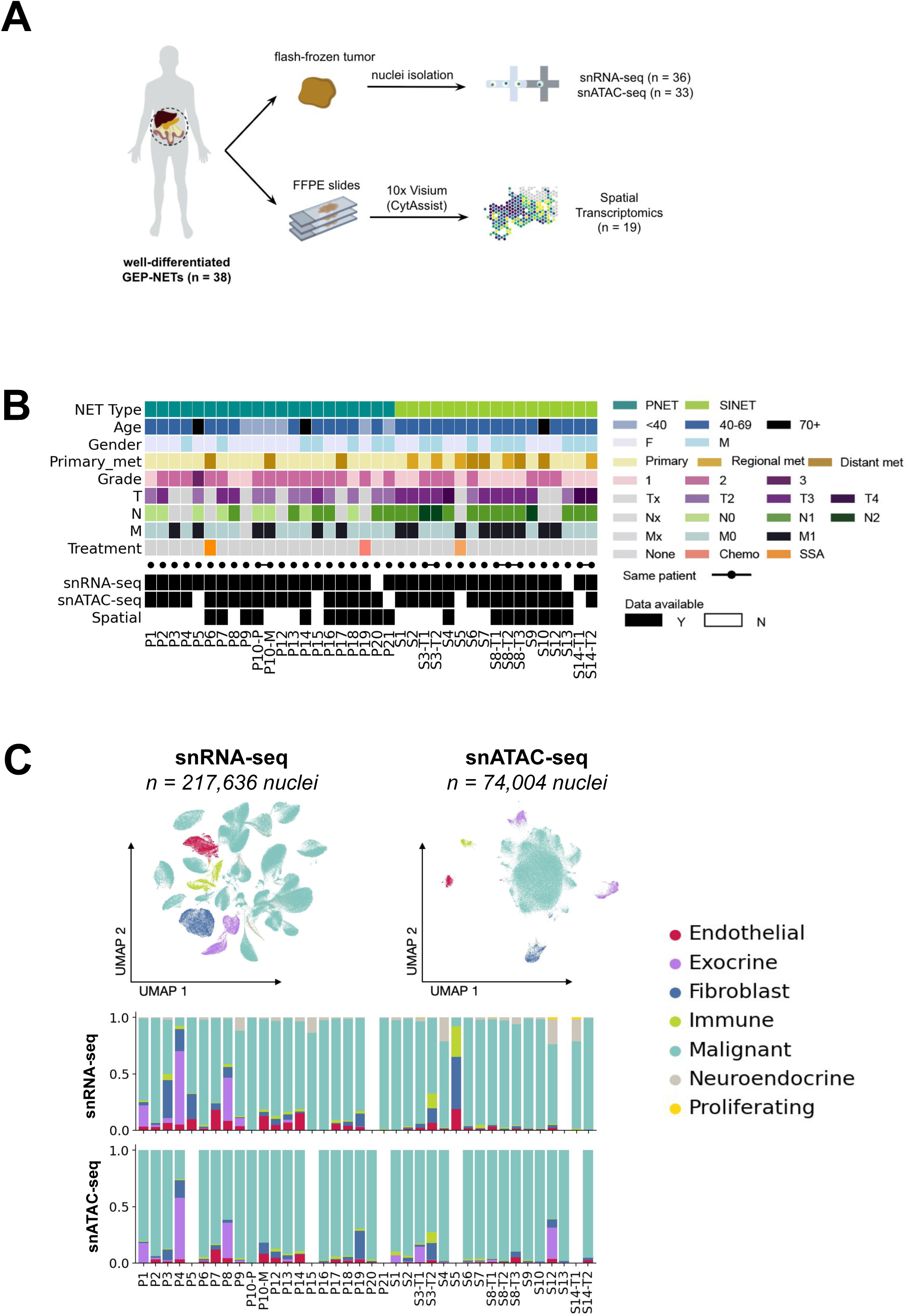
Multiomic characterization of well-differentiated pancreatic and small intestinal neuroendocrine tumors. (A) Multiome sample processing and sequencing workflow. Archival, flash-frozen NET samples were apportioned for nuclei isolation and subsequent joint single-nuclei RNA-sequencing and ATAC-sequencing using the 10x Genomics protocol. Formalin-fixed, paraffin-embedded (FFPE) slides were generated from each sample and used to generate spatial transcriptomics data using the 10x Genomics Visium protocol. (B) Clinical characteristics of the cohort, including tumor origin, stage, grade, treatment status, and availability of multi-omic data after quality control. (C) UMAP embeddings of snRNA-seq and snATAC-seq data colored by annotated cell types. Bar plots indicate the relative abundance of major cell populations across samples.

25 of the samples in our cohort were primary tumors resected from either the pancreas or small intestine. Of the remaining samples, 5 were regional metastatic siNETs, and 7 were distant liver metastases originating from either pNETs (n = 3) or siNETs (n = 4) (Figure 1B, Supplementary Table 1). At the time of tumor resection, all but three of the NETs in our cohort were treatment-naive (Figure 1B). Among the three with prior therapy exposure, two patients had received somatostatin analog therapy, and one pNET patient had received combination capecitabine/temozolomide therapy. Of all the samples in the cohort, one pNET was classified as World Health Organization Grade 3 (WHO G3); the remaining tumors were classified as WHO G1 or G2 (Figure 1B).

Next, we defined the cellular landscapes of pNETs and siNETs captured within our multiomic single-nuclei sequencing data. After quality control and pre-processing, our snRNA-seq dataset included 217,636 high-quality nuclei from 36 of our 38 samples (Figure 1C, Methods). Leiden clustering and canonical marker gene expression were used to define broad cell types, including immune, endothelial, cancer-associated fibroblast (CAF), exocrine, and epithelial (Figure 1C, Supplementary Figure 1A). Then, we combined inferred copy number alterations (CNAs) with expression of established NET tumor markers to distinguish malignant NET cells from non-malignant populations (Methods, Supplementary Figure 1B). We also obtained 74,004 single-nuclei chromatin accessibility profiles from 33 of our 38 samples (Figure 1C) after quality control and filtering. Broad cell types were assigned using label transfer from matched snRNA-seq barcodes and clustering (Methods). Malignant cells composed the largest proportion of cell types recovered in both snRNA-seq and snATAC-seq (Figure 1C). Among microenvironmental populations, CAFs were the most frequently recovered cell types, followed by normal exocrine cells, endothelial cell types, and immune cell populations (Figure 1C).

### GEP-NET malignant cells comprise a spectrum of neuroendocrine transcriptional phenotypes

Well-differentiated GEP-NETs are currently classified based on anatomical site and histologic grade. However, this classification does not account for tumor-intrinsic transcriptional heterogeneity, nor does it reflect potential differences in cell states that may influence behavior or therapeutic response. Genomic studies have identified recurrent alterations in *MEN1*, *ATRX*, *DAXX,* and mTOR pathway genes in pNETs (9–11), while siNETs typically harbor few recurrent somatic driver mutations (12). Yet, the extent to which transcriptional and epigenetic programs shape conserved and divergent tumor phenotypes across NETs remains unresolved.

To gain understanding of the transcriptional and epigenetic programs within NET cells, we isolated the malignant cells and applied consensus non-negative matrix factorization (cNMF) separately to siNET and pNET malignant cells to uncover recurrent transcriptional programs (Methods, Figure 2A-B). In siNETs (n = 76,010 tumor cells from 16 tumors), we identified three robust cNMF programs (Figure 2A, Supplementary Table 2). The first program (si-cNMF1) was enriched for neuronal and synaptic signaling genes, including *KCNJ3, RBFOX1, DLG2, NRXN1,* and *CADPS,* as well as calcium channel subunits (*CACNA2D1*) and neurodevelopmental transcription factors (TFs; *SOX5, DAB1*). Through gene set enrichment analysis (GSEA), we identified that this program was enriched for pathways involved in apical junction, mitotic spindle, and L-2/STAT5 signaling (Figure 2C). The second program (si-cNMF2) reflected secretory neuroendocrine identity, marked by top associated genes *CHGA, CHGB, PCSK1, SCG5,* and *TAC1*, and was enriched for oxidative phosphorylation, adipogenesis, and mTORC1 signaling pathways. The third (si-cNMF3) displayed features of epithelial-to-mesenchymal transition (EMT) and tumor progression, with top associated genes *ZEB2, ROBO1, DOCK4,* and *AHNAK*, along with stem-like (*TCF4*) and stress-responsive genes (*LGALS3, NEAT1*).

**Figure 2:**
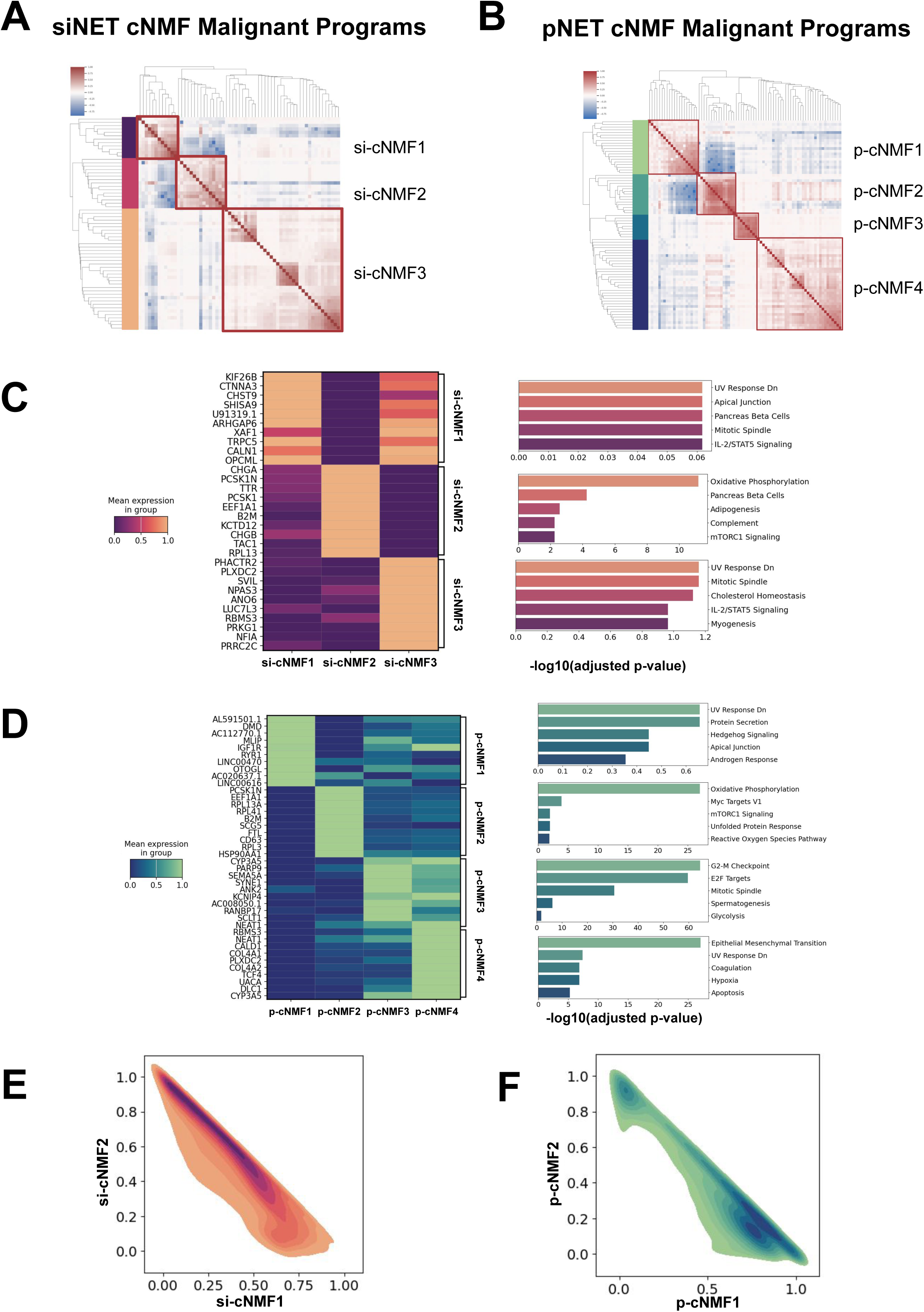
Identification and characterization of malignant cNMF programs in siNETs and pNETs. (A,B) Heatmaps showing gene weights for cNMF programs identified independently in malignant cells from siNETs (A) and pNETs (B). (C,D) Gene set enrichment analysis (GSEA) of hallmark pathways associated with each cNMF program. Normalized enrichment scores (NES) and false discovery rate (FDR)–adjusted q-values are shown. (E,F) Two-dimensional density plots of normalized neuronal-like (cNMF1) versus secretory neuroendocrine (cNMF2) program scores in siNETs (E) and pNETs (F), illustrating a continuous transcriptional spectrum.

Similarly, we identified four cNMF programs in pNET tumor cells (n = 100,809 tumor cells from 20 tumors; Figure 2B, Supplementary Table 3). These included a neuronal-like program (p-cNMF1) with top associated genes *RIMS2, NRXN1, MEIS2, FGF14, NBEA*, and TFs *NPAS3, TFAP4*, and *ISL1*. This program was enriched for protein secretion, hedgehog signaling, and apical junction pathways (Figure 2D). The secretory neuroendocrine program (p-cNMF2) mirrored that seen in siNETs, with expression of *CHGA, CHGB, SCG2–5, TTR*, and *CPE*, and enrichment for oxidative phosphorylation, MYC targets, and mTORC1 signaling pathways. A third program (p-cNMF3) corresponded to a proliferative state with top associated genes *MKI67, CDK1, CCNB1, AURKA, BIRC5, CDC20*, and *TOP2A*, and was enriched for G2/M checkpoint and E2F targets. The final program (p-cNMF4) represented an EMT and stromal state defined by *VIM, FN1, COL1A1, COL3A1, SPARC*, and other matrix remodeling genes.

To determine whether these cNMF programs reflected shared cellular states across NET types, we also performed integrated cNMF on pooled siNET and pNET tumor cells. This analysis recapitulated the major cNMF programs identified independently as outlined above, and, importantly, demonstrated that each program included cells from both tumor types (Supplementary Figure 2). Two-dimensional density plots of normalized program scores (neuronal-like si/p-cNMF1 vs. secretory neuroendocrine si/p-cNMF2) demonstrated that respective malignant siNET or pNET cells form a continuous distribution across the two programs (cNMF1/2) rather than resolving into distinct subtypes (Figure 2E-F). These findings support a model in which, regardless of the organ site, GEP-NET malignant cells span a conserved transcriptional spectrum anchored by neuronal-like and secretory neuroendocrine identities.

### Divergent Epigenetic Landscapes Drive Organ Site-Specific Tumor Programs

We next asked whether this conserved transcriptional continuum among the two predominant programs was regulated by shared or organ site-specific epigenetic programs. We leveraged matched ATAC-seq data and performed TF binding motif enrichment analysis on accessible chromatin regions associated with each cNMF program, stratified by originating organ type (Methods). Despite transcriptional convergence with respect to the cNMF program enrichments, siNETs and pNETs displayed distinct epigenetic regulatory signatures. In siNETs, malignant cells most enriched for the the neuronal-like cNMF program (si-cNMF1) had accessibility around binding motifs for TFs *MAX::MYC*, *MITF*, and circadian regulators such as *CLOCK* and *TP63*, suggesting a stress-responsive, neurodevelopmental regulatory network similar to sensory or peripheral neurons (13,14) (Figure 3A). The secretory program (si-cNMF2) was enriched around binding motifs for TFs *HOXA9*, *JUN::FOS*, and *SOX9*, which are linked to lineage plasticity, stem-like states, and epithelial remodeling (15,16) (Figure 3B). When examining the effect on gene expression, we observed that the enriched TF binding motifs were associated with increased expression of their predicted target genes (Figure 3C). This observation reinforces the link between regulatory motif accessibility and downstream transcriptional expression, illustrating how distinct TF networks shape the neuronal-like versus secretory cNMF programs.

**Figure 3:**
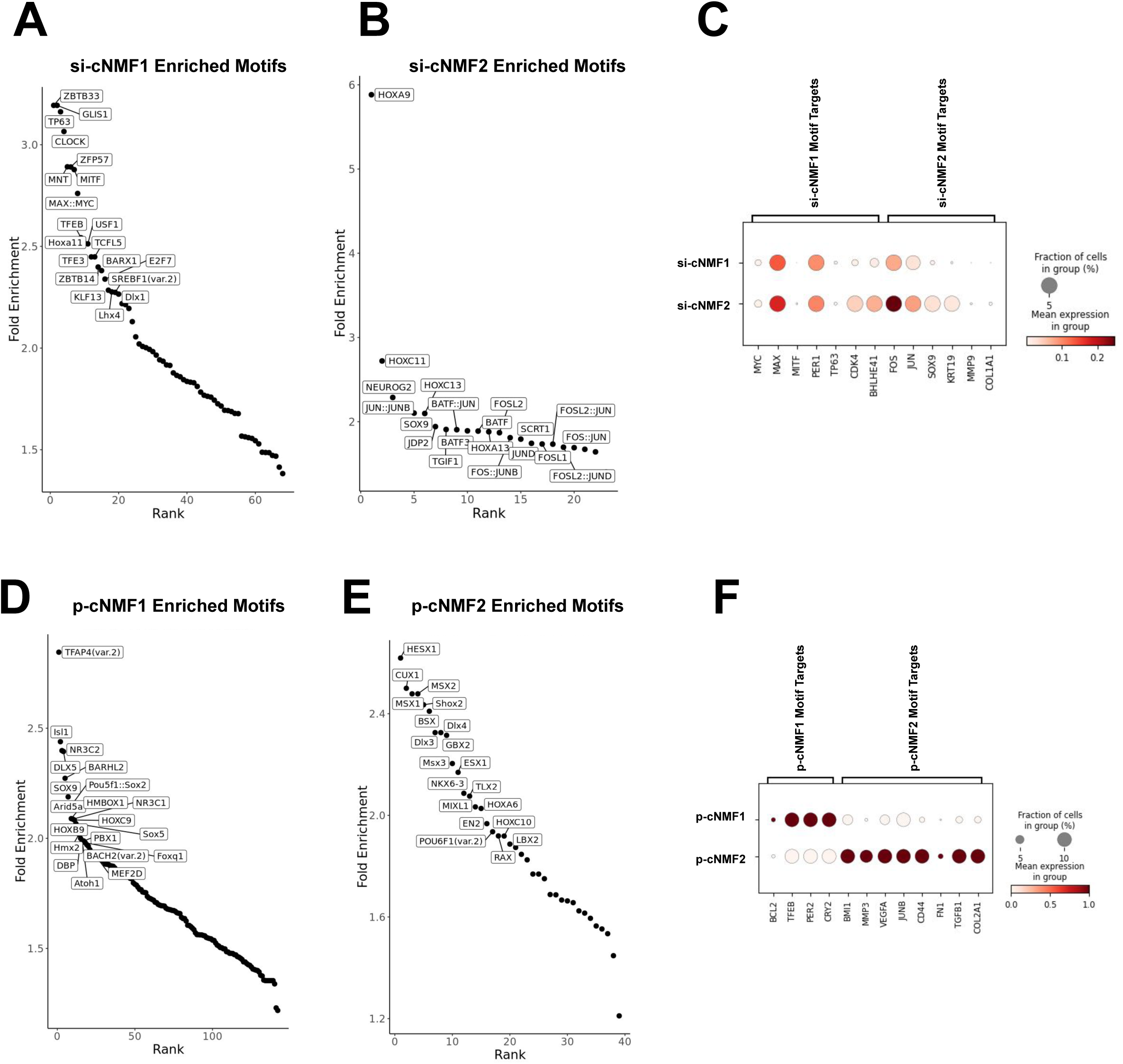
Distinct epigenetic landscapes regulate conserved malignant transcriptional programs in siNETs and pNETs. (A,B) Transcription factor (TF) binding motif enrichment analysis of snATAC-seq peaks associated with siNET malignant cells enriched for the neuronal-like cNMF1 program (A) and secretory cNMF2 program (B). Motif enrichment significance was assessed using chromVAR deviation scores. (C) Expression of predicted target genes corresponding to enriched TF motifs in siNET malignant cells. (D,E) TF motif enrichment for pNET malignant cells enriched for neuronal-like p-cNMF1 (D) and secretory p-cNMF2 (E) programs. (F) Expression of predicted target genes for pNET-enriched motifs. Comparisons are descriptive; statistical testing was performed as described in Methods.

By contrast, in pNETs, the malignant cells most enriched for the p-cNMF1 program displayed chromatin accessibility around binding motifs for TFs *TFAP4*, *ISL1*, and *NR3C2*, TFs involved in foregut and pancreatic development (17,18) (Figure 3D). pNET malignant cells most enriched for the secretory neuroendocrine program (p-cNMF2) displayed accessibility around hindbrain and midbrain-associated TF binding motifs, including *MSX2*, *LBX2*, and *EN2*, suggesting an alternate developmental origin for regulatory control (19–21) (Figure 3E). Importantly, these TF binding motif enrichments were largely non-overlapping between siNETs and pNETs despite the similarity in associated cNMF programs, indicating that the convergent transcriptional programs arise within divergent epigenetic frameworks shaped by tissue of origin. Again, we observed that predicted gene targets of the enriched TF regulons had higher expression (Figure 3F).

These findings suggest that while siNETs and pNETs converge on a common axis of transcriptional malignant cell programs, spanning neuronal-like to secretory neuroendocrine, the upstream epigenetic regulatory circuits associated with these malignant cell programs are organ-specific. Notably, the secretory neuroendocrine program in both tumor types, which is defined by classic neuroendocrine identity, was also enriched for regulatory motifs associated with developmental plasticity and migration, raising the possibility that this transcriptional state may be preferentially selected or stabilized within organ-specific epigenetic contexts during tumor progression.

### Spatial coupling of malignant programs with distinct microenvironmental niches

To further understand how these GEP-NET transcriptional programs are shaped by and influence the tumor microenvironment, we next assessed the spatial organization of the usage of these transcriptional programs using 10x Visium spatial transcriptomics (ST). For these samples, we categorized ST spots as pure tumor regions, mixed regions containing both malignant cells and TME cells in similar proportions, or regions of normal tissue, using CNA profiles inferred from the spatial data (Methods).

Consistent with pathologist annotation and similar to the single-nucleus multiome cell data, the spatial transcriptomics data had a high prevalence of malignant cells (Figure 4A). We next performed gene signature scoring for the three siNET and four pNET malignant cell cNMF programs on the inferred tumor spots and found that they displayed distinct spatial distributions within the GEP-NET samples (Methods). To further characterize spatial organization, we conducted distance-based analyses to evaluate the localization of malignant spots enriched for each cNMF program relative to the tumor boundary, as well as their proximity to neighboring cell types inferred from adjacent spots (Methods). Notably, malignant spots with high cNMF2 expression were preferentially localized deeper within the tumor interior, whereas cNMF1-high malignant spots were more frequently found closer to the tumor periphery (Figure 4B), indicating that conserved malignant transcriptional programs occupy distinct spatial niches within GEP-NET tumors.

**Figure 4:**
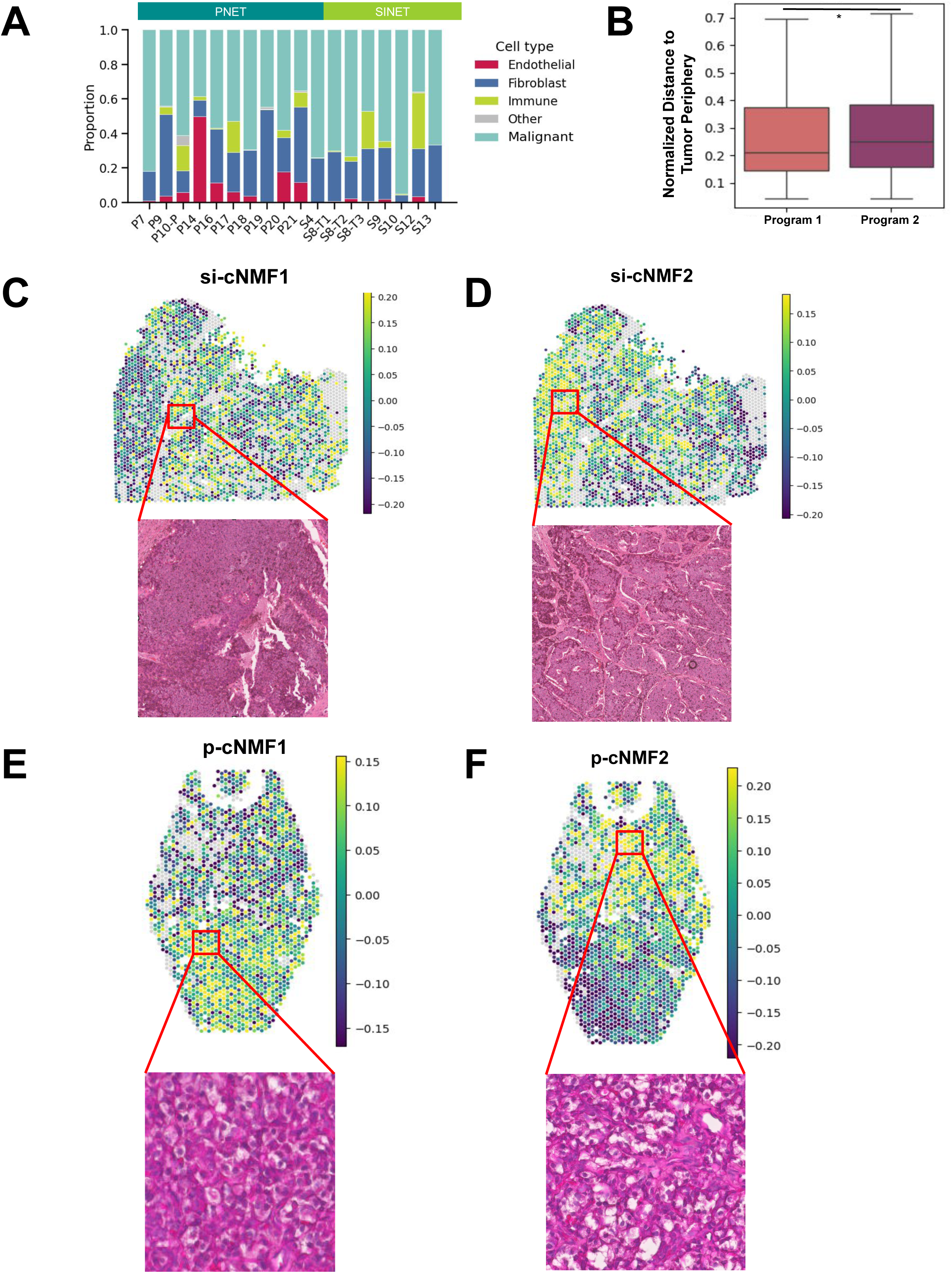
Spatial transcriptomics reveals distinct localization of malignant cNMF programs within tumors. (A) Proportions of inferred malignant, stromal, immune, and normal tissue regions across spatial transcriptomic samples. (B) Distance of malignant spots enriched for cNMF1 or cNMF2 from the tumor boundary. Distances were compared using a two-sided Wilcoxon rank-sum test. (C,D) Representative siNET spatial maps showing enrichment of cNMF1-high malignant spots in densely cellular regions (C) and cNMF2-high malignant spots in stromal-rich regions (D). (E,F) Spatial distribution of pNET malignant spots enriched for p-cNMF1 (E) and p-cNMF2 (F). All spatial maps are representative examples.

When examining the histology of the areas enriched for the cNMF programs, we found that malignant spots scoring highest for si-cNMF1 were specifically localized in areas with high density of tumor cells, while malignant spots scoring highest for si-cNMF2 were present in areas with stromal infiltration (Figure 4C and 4D, Supplementary Figure 3A-D). For pNET samples, we likewise observed spatial localization of the malignant spots scoring highest for the respective cNMF programs, where malignant spots scoring most highly for p-cNMF1 were positioned in areas with high tumor cell density, and malignant spots scoring most highly for p-cNMF2 comprised an area of cells with degraded cytoplasms (Figure 4E and 4F).

Transcriptional profiles of the spots confirmed these spatial patterns. Malignant spots scoring highest for the si/p-cNMF2 program showed significantly increased expression of genes involved in extracellular matrix remodeling, including *COL1A1*, *SPARC*, and *MMP2* (FDR-adjusted p = 0.015, paired Wilcoxon test across samples), whereas malignant spots scoring highest for the si/p-cNMF1 program exhibited strong enrichment of antigen presentation–associated genes (FDR-adjusted p = 1.9 × 10⁻⁵). Consistent with these transcriptional differences, si/p-cNMF1-enriched malignant spots localized to immune-dense tumor regions, whereas si/p-cNMF2-enriched malignant spots preferentially occupied stromal- and vascular-associated niches. Together, these spatial associations suggest that conserved malignant transcriptional programs manifest within distinct tumor-microenvironmental niches.

### si/p-cNMF2 defines a vascular-interactive malignant program primed for stromal remodeling and metastatic dissemination

GEP-NETs have uniquely high vascularization among solid tumors, a feature that underpins their pathophysiology and positive clinical responses to anti-angiogenic therapies such as sunitinib, bevacizumab, surufatinib, and cabozantinib (13,22–25). However, the NET phenotypes that drive vascular remodeling and their spatial coordination with stromal compartments remain incompletely defined.

To investigate how the identified malignant cNMF programs may influence how the malignant cells coordinate with other microenvironmental cell types to support NET invasion and progression, we performed ligand-receptor interaction analysis within the snRNA-seq data (Methods). Across both siNET and pNET datasets, si/p-cNMF2-high malignant cells exhibited enriched paracrine signaling interactions with endothelial cells and fibroblasts, particularly through pathways known to mediate angiogenesis, fibrosis, and extracellular matrix remodeling (Supplementary Figure 4). Top-ranked interactions included *TGFB1-ITGB1*, *VEGFA-FLT1*, and *LAMA2-ITGA1*, suggesting si/p-cNMF2-high malignant cells may be engaged in the activation of perivascular and stromal niches (26–28).

We next explored how these interactions might relate to the spatial context with our spatial transcriptomic data. Across all samples, si/p-cNMF2-high malignant spots were significantly more often adjacent to endothelial cells than si/p-cNMF1-high malignant spots (p < 0.05; Wilcoxon two-sided test) (Figure 5A). For example, in a representative pNET sample (P17), endothelial cells were consistently localized next to cNMF2-high (secretory) malignant spots (Figure 5B-C). These regions exhibited elevated expression of pro-fibrotic (*TGFB1, ANGPT1*), matrix-remodeling (*MMP2, COL1A1*), and angiogenic genes (*VEGFA, SPARC*) (p = 8.23 × 10⁻³; Wilcoxon rank-sum test) (Figure 5D). Moreover, endothelial cells in these regions expressed high levels of *FLT1, TEK,* and *TGFBR2*, supporting active paracrine signaling with adjacent tumor cells.

**Figure 5.**
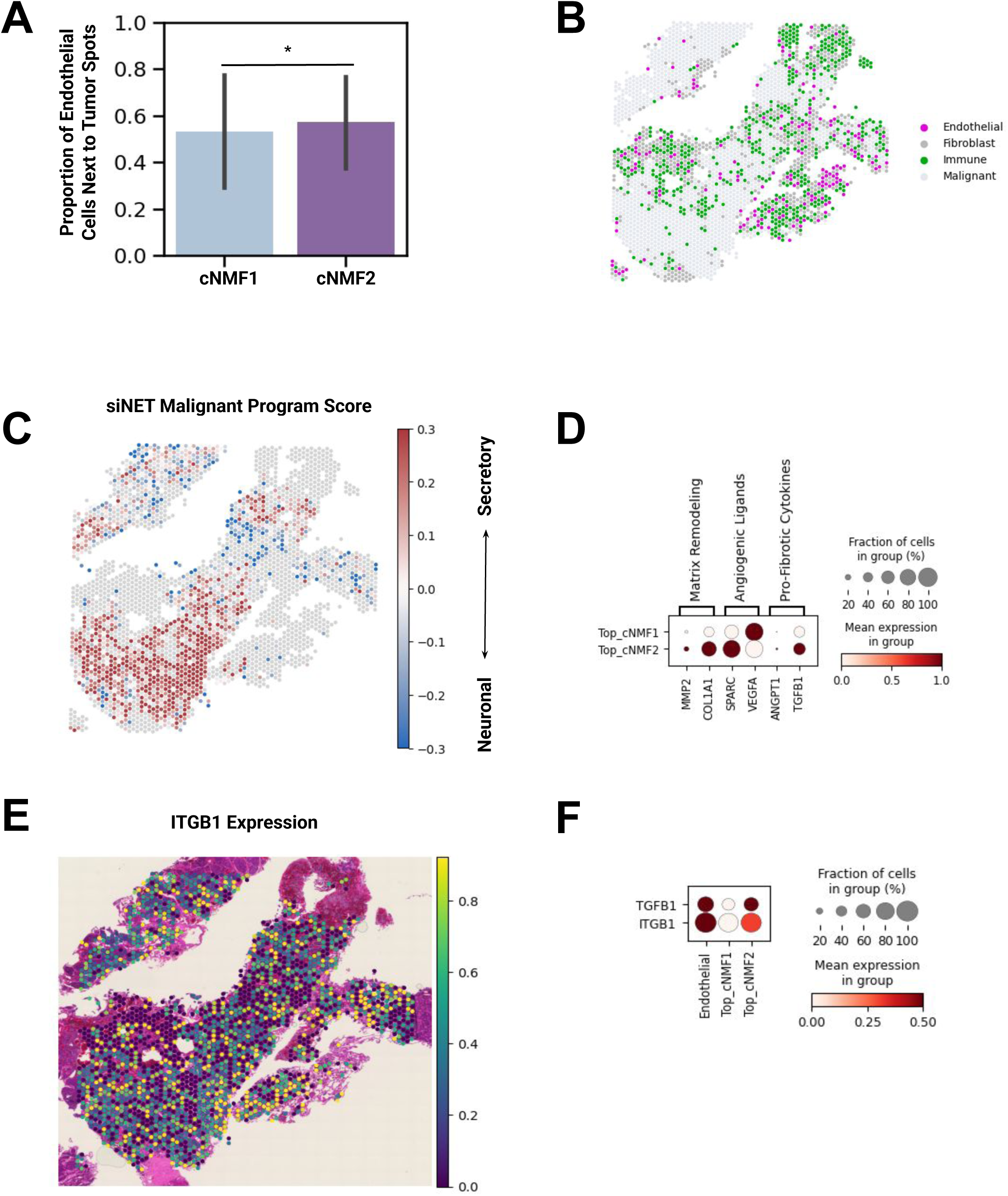
si/p-cNMF2-high malignant cells spatially associate with endothelial cells and exhibit stromal and angiogenic transcriptional programs. (A) Fraction of endothelial cells adjacent to cNMF1-high versus cNMF2-high malignant spots across spatial transcriptomic samples. Statistical significance was assessed using a two-sided Wilcoxon signed-rank test across samples. (B,C) Representative pNET spatial maps showing localization of endothelial cells adjacent to cNMF2-high malignant spots. (D) Dot plot showing expression of pro-fibrotic, extracellular matrix remodeling, and angiogenic genes in malignant spots stratified by cNMF program. Per-gene expression differences were assessed using a Wilcoxon rank-sum test, with p-values adjusted for multiple testing using the Benjamini–Hochberg method. (E,F) Spatial expression of TGFB1 and ITGB1, demonstrating polarized expression in cNMF2-high malignant regions.

Relatedly, the ligand-receptor interaction analysis within the snRNA-seq data predicted tumor-endothelial communication, with conserved ligand-receptor pairs that promote neovascularization and vascular permeability (Supplementary Figure 4) (29). Notably, *TGFB1* expression was polarized, with high expression in si/p-cNMF2-high malignant spots but with negligible expression in si/p-cNMF1-high malignant spots, paralleling *ITGB1* expression patterns (Figure 5E-F). In parallel, TF binding motif enrichment analysis of the snATAC-seq data revealed increased accessibility at *AP-1* (*JUN::FOS*) and *SMAD* motifs in si/p-cNMF2-high malignant cells, transcription factors that act as downstream effectors of cellular stress and TGFβ signaling pathways. Given the established roles of *AP-1* and TGFβ signaling in extracellular matrix remodeling and stromal-tumor interactions (26,30), these findings are consistent with a chromatin landscape that may be permissive to stromal engagement in cNMF2-high malignant cells, rather than indicative of active regulatory control.

Together, these findings position the si/p-cNMF2 malignant cell program as the dominant invasive program in GEP-NETs, responsible for spatially organized vascular co-option and stromal remodeling. The spatial and molecular features of the si/p-cNMF2 program may explain the existing clinical efficacy of anti-angiogenic therapies in GEP-NETs and further highlight the tumor–endothelial interface as a therapeutic target.

### si/p-cNMF1 represents a neuronal-like, immune-engaged tumor state with therapeutic implications

In contrast to the stromal-interactive phenotype of malignant cell program si/p-cNMF2, si/p-cNMF1-high malignant cells had predicted interactions with myeloid populations, particularly macrophages and monocytes. The described ligand-receptor interaction analysis revealed conserved neuroimmune signaling across si/p-cNMF1-high malignant cells, including *CX3CL1-CX3CR1*, *IL34-CSF1R*, and *CSF1-CSF1R*, pathways known to regulate microglial-neuronal crosstalk, monocyte recruitment, and innate immune niche formation (31–34) (Supplementary Figure 4). Additionally, si/p-cNMF1-high malignant cells expressed neuronal ligands such as *NRXN1, DLG2*, and *RIMS2*, highlighting enrichment of neuronal communication-related genes within immune-interactive malignant regions (35,36) (Supplementary Figure 5).

The spatial transcriptomic data supported this myeloid-engaged state. Malignant spots scoring highest for the si/p-cNMF1 program preferentially localized to densely cellular tumor regions and were consistently found in proximity to myeloid cell populations in both siNETs (Figure 6A) and pNETs (Figure 6B). In a representative siNET (sample S4), si-cNMF1-high malignant spots (neuronal) were positioned within an immune-dense cluster (Figure 6C and 6E). The malignant spots in these immune-enriched regions lacked enrichment of si-cNMF2 and these regions were notably devoid of fibrotic or vascular features, reinforcing a distinct immune-permissive microenvironment. By contrast, pNETs, characteristically more vascularized, exhibited higher immune infiltration overall, yet still displayed a reciprocal pattern where p-cNMF2-high malignant spots were present in regions of low immune density, as observed in sample P17 (Figure 6D and 6F).

**Figure 6.**
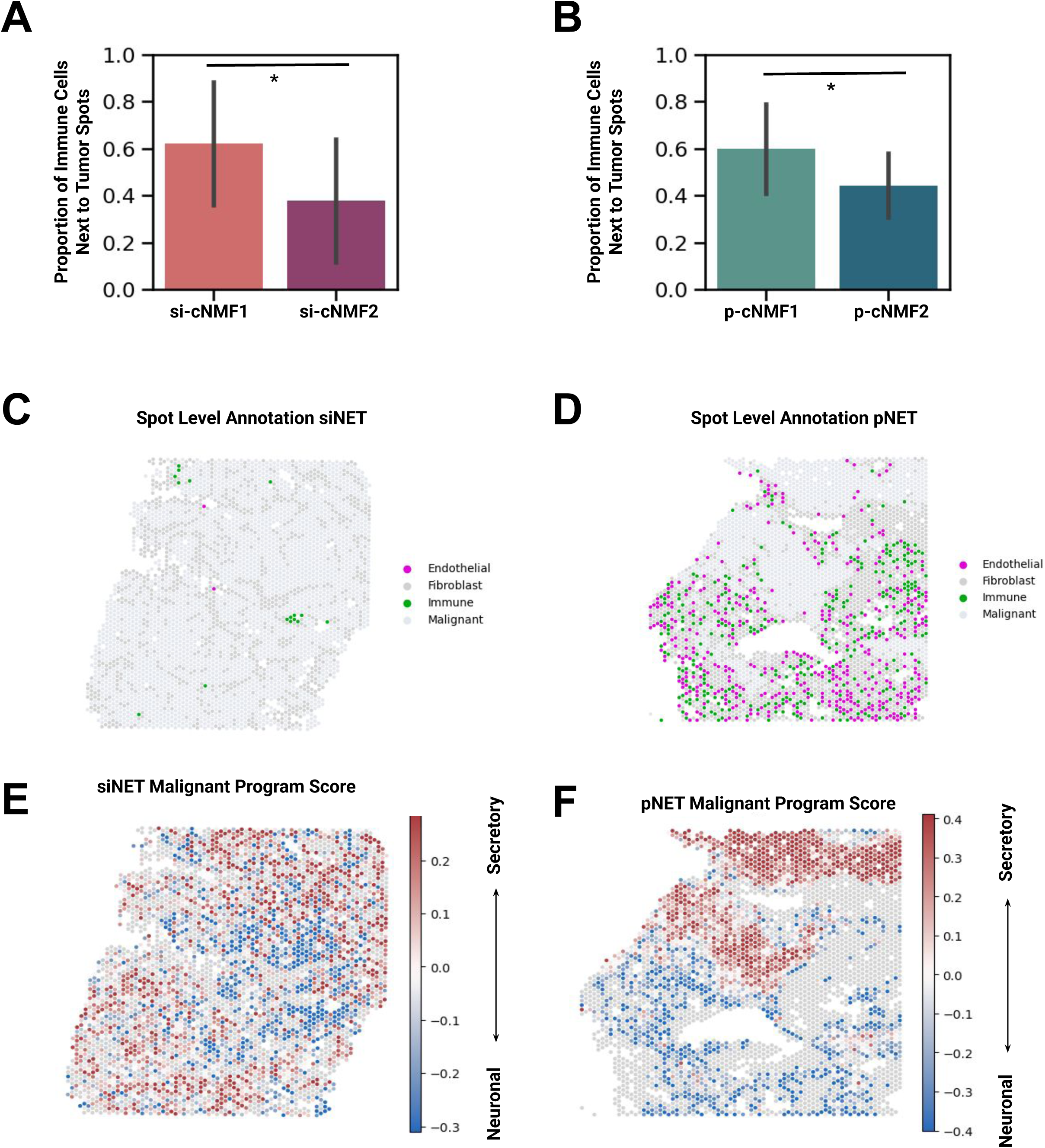
Neuronal-like cNMF1-high malignant cells localize to immune-rich tumor regions. (A,B) Proportion of immune cells adjacent to cNMF1-high versus cNMF2-high malignant spots in siNETs (A) and pNETs (B). Differences were assessed using a two-sided Wilcoxon signed-rank test across samples. (C,D) Representative spatial maps showing immune cell localization relative to malignant spots in siNET (C) and pNET (D) samples. (E,F) Spatial distribution of malignant cNMF program scores highlighting immune-dense regions enriched for cNMF1-high malignant spots.

These findings suggest that neuronal-like si/p-cNMF1-high malignant cells actively recruit and modulate myeloid cells by co-opting neuroimmune signaling programs, forming spatially confined, myeloid-rich niches that support tumor growth through the aforementioned signaling interactions. This myeloid-engaged phenotype was most pronounced in siNETs, and was distinct from the stromal- and vascular-dominant landscapes associated with the secretory neuroendocrine-like si/p-cNMF2-high malignant cells.

Overall, the neuronal-like si/p-cNMF1 malignant cells represent a neuroimmune-interacting state, sharply contrasted with the stromal- and angiogenesis-driven secretory neuroendocrine malignant cell program, si/p-cNMF2. The divergent spatial ecosystems associated with the malignant cells enriched for the respective cNMF programs raise the possibility that effective treatment of GEP-NETs could involve strategies tailored to the transcriptional and microenvironmental context of the different cell programs.

### cNMF programs associate with NET clinical stage and disease progression

Lastly, we evaluated how the malignant cNMF programs relate to clinical stage and tumor progression in both siNET and pNET tumors. Across both tumor types, si/p-cNMF1 was predominantly enriched in localized primary tumors, whereas si/p-cNMF2 was significantly enriched in metastatic and recurrent lesions (Figures 7A and 7B).

**Figure 7.**
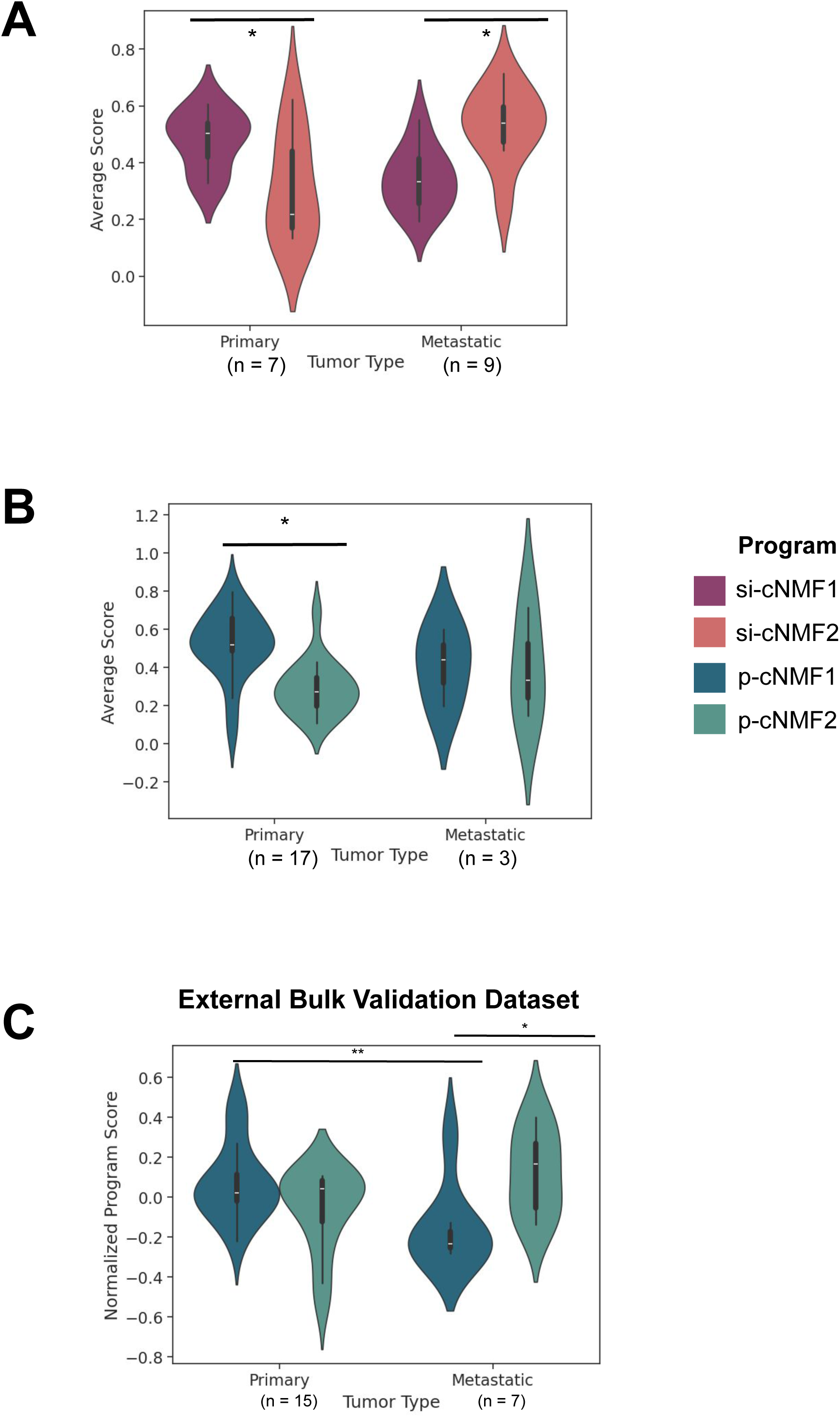
Malignant transcriptional programs associate with clinical stage and metastatic progression. (A,B) Violin plots showing cNMF program scores stratified by tumor type and clinical stage for siNETs (A) and pNETs (B). Differences between primary and metastatic tumors were assessed using a two-sided Wilcoxon rank-sum test. (C) Validation in an independent bulk RNA-seq pNET cohort showing decreased neuronal p-cNMF1 scores and increased secretory p-cNMF2 scores in metastatic samples. Statistical significance was assessed using a two-sided Wilcoxon rank-sum test (* = p value < 0.05, ** = p value < 0.01).

In siNETs, this shift was most pronounced along the metastatic continuum, with si-cNMF2 scores increasing from regional to distant metastatic disease. Consistent with this observation, malignant cNMF program scores were also associated with metastatic status at diagnosis, with higher si/p-cNMF2 and lower si/p-cNMF1 scores observed in patients presenting with metastatic disease (Supplementary Figure 5). These patterns were reproducible within individual tumors sampled across progression states, including a representative siNET sample (S8) that exhibited a stepwise increase in si-cNMF2 and concomitant decrease in si-cNMF1 from primary to regional and distant metastatic lesions (Supplementary Figure 5).

To further validate the association between malignant cell programs and tumor progression, we analyzed an independent external bulk RNA-seq pNET dataset (37). In this independent dataset, metastatic tumors exhibited significantly reduced neuronal p-cNMF1 scores and significantly elevated secretory neuroendocrine p-cNMF2 scores compared to primary tumors (Figure 7C). Together, these findings demonstrate that these malignant cNMF programs exhibit distinct clinical associations with primary and metastatic NETs.

## DISCUSSION

Here, we present the largest and most comprehensive single-nucleus and spatial transcriptional profiling study of malignant cells in GEP-NETs to date. We integrated single-nucleus multi-omic and spatial transcriptomic data to reveal the transcriptional, epigenetic, and spatial features that shape NET heterogeneity. To our knowledge, this study is the first to place malignant cell states in their spatial context across GEP-NETs, allowing us to spatially associate the identified malignant cell programs and the niche-specific tumor-microenvironment interactions predicted that drive NET growth and progression.

Altogether, our analyses identified two conserved malignant cell cNMF programs that structure heterogeneity across pNETs and siNETs. Despite their distinct genomic landscapes as described in prior studies (6), we found that tumors from both organ sites converge on neuronal-like (si-cNMF1/p-cNMF1) and secretory neuroendocrine (si-cNMF2/p-cNMF2) programs, underscoring a shared architecture of NET biology.

We further discovered that these programs are active, functional states characterized by distinct epigenetic regulation, spatial localization, and clinical associations.

The secretory si/p-cNMF2 program was enriched in stromal-vascular niches, where it engaged extracellular matrix remodeling and angiogenic pathways. Its spatial localization and enrichment in metastatic tumors suggest that si/p-cNMF2 represents a metastasis-enabling state, potentially driven by TGFB-integrin crosstalk and other stromal interactions. In contrast, the neuronal si/p-cNMF1 program localized to high cell density, immune-interactive regions and associated with calcium signaling and neuronal TF motifs. This program was enriched in non-metastatic tumors, consistent with a more plastic, progenitor-like phenotype capable of immune engagement. Together, these findings support a model in which GEP-NET progression is associated with a shift in the relative prevalence of transcriptional programs, with advanced and metastatic tumors preferentially enriched for the secretory si/p-cNMF2 program. The neuronal si/p-cNMF1 program, with lineage-restricted TF (e.g., *TFAP4*, *MITF*, *Isl1*) binding motif enrichment, may represent a more differentiated, tissue-committed program. By contrast, si/p-cNMF2, marked by *HOX*, *AP-1*, and developmental TF binding motif enrichment, displays hallmarks of dedifferentiation and plasticity, enabling cells to remodel their identity and adapt to stress. This dichotomy mirrors paradigms in other solid tumors, where conserved and transient transcriptional states provide a more powerful framework for understanding disease progression than mutational profiles alone (38).

Our results also extend prior NET classifications based on genotype or proliferation index by highlighting a unifying cell-state model that transcends the originating organ site. Importantly, we identify putatively actionable signaling pathways within these states, including TGFB-integrin crosstalk in the secretory si/p-cNMF2 program and myeloid-tumor interactions in the neuronal si/p-cNMF1 program. Therapeutically, such features open potential opportunities for intervention. These include myeloid-directed therapeutics currently in clinical development that could be used to disrupt tumor-myeloid interactions in neuronal-like NETs. The spatial confinement of these niches may also enable localized delivery of immunomodulatory agents while sparing stromal and vascular compartments. These insights open avenues for therapeutic strategies that target tumor-microenvironment interactions or exploit plasticity in NET differentiation states.

Limitations to this study include the relatively modest number of spatial transcriptomic samples and the need for longitudinal data to fully define the dynamics of program switching during progression or therapy. We were also limited in the spatial resolution of the data in this study. Emerging spatial technologies, such as Visium HD or Xenium, may provide higher resolution information about the cell types. This approach would be especially helpful in this broadly malignant cell-dominant solid tumor type and the related interactions within the microenvironment. Additionally, the lack of robust *in vitro* and *in vivo* preclinical models for NETs limits the ability to functionally interrogate the findings reported herein. Once such models become available, future studies should investigate the mechanisms and potential therapeutic targets nominated in our work in the *in vitro* setting.

In summary, our study establishes a cross-tissue framework for understanding NET biology through conserved malignant cell programs. By linking these programs to spatial niches, microenvironmental interactions, and clinical stage, we provide a roadmap for tumor progression and therapeutic targeting in GEP-NETs.

## METHODS

### Lead Contact

Further information and other requests should be directed to the corresponding author.

### Data and Code Availability

Raw single-nuclei RNA-seq and single-nuclei ATAC-seq data will be deposited into dbGaP (accession number phs003141.v2). Processed single-nuclei RNA-seq data will be deposited in the Broad Institute Single Cell Portal (https://singlecell.broadinstitute.org/single_cell). Processed spatial data will be deposited in Zonedo. Code to reproduce all figures will be made available in a GitHub repository.

### Institutional Approval

Human patient samples were collected under Dana-Farber Cancer Institute IRB protocol #02-314. Informed consent was required for patient participation, which was obtained per protocol: At the time of their clinic appointments, patients were contacted by a member of the study staff and were asked if they were interested in participating. The study was explained in detail, and informed consent was signed if the patient/control ultimately wished to participate. Patients were specifically consented to 1) allow the collection of tissue and blood 2) to allow the collection and linkage of clinical data to the tissue 3) to allow for re-contact of the patient for the purpose of recruitment into new clinical protocols, or for the purpose of obtaining clinical follow-up, 4) to allow for a one-time additional blood draw or urine collection for study purposes if not feasible in the context of a scheduled blood draw, and 5) collection of periodic blood samples.

## Experimental Methods

### Sample collection and nuclei isolation for single-nuclei RNA-seq/ATAC-seq

Fresh tumor resection tissue was collected from consented patients at Brigham and Women’s Hospital (BWH). Multiple pathologists in the BWH Department of Surgical Pathology identified tumor tissue within each resection, which was dissected away from normal tissue using a scalpel. Tumor tissue that was not needed for clinical pathology review was snap-frozen and held in liquid nitrogen tanks for long-term storage.

To isolate nuclei for joint multiomic single-cell RNA-seq/ATAC-seq, a portion of each snap-frozen tumor sample was cut off with a scalpel and placed into 1.5mL Eppendorf tubes on dry ice. 1mL of TST buffer was then added to the tube. Tumor samples were minced inside the tube into <0.5mm fragments with spring scissors for seven minutes on ice, followed by three minutes of mechanical dissociation by pipetting the tumor-buffer mixture up and down using a P1000 tip. At the end of the 10-minute mincing period, the contents of the tube were transferred to a 30μm filter (MACS SmartStrainers, Miltenyi #130-098-458) attached to a 15mL Falcon tube on ice. An additional 1mL of TST was used to rinse the Eppendorf tube and the 30μm filter. The filter was then rinsed with 3mL of 1X TST buffer, bringing the total suspension volume to 5mL.

Next, the suspension was centrifuged in a swinging bucket rotor for 5 minutes at 500xg at 4°C with the centrifuge deceleration ramp set to 5 (out of maximum 9). The 15mL tube containing the cell pellet was placed on ice, and the supernatant was carefully removed using P1000 and P200 pipettes. To lyse cells, 100μL of 0.1X Lysis Buffer was added to the pellet, which was gently dissociated by pipetting 5 times with a P200 pipette. After two minutes, 1mL of Wash Buffer was added and gently pipetted 5 times to prevent excessive lysis. The resulting suspension was centrifuged in a swinging bucket rotor for 5 minutes at 500xg at 4°C with the centrifuge deceleration ramp set to 5.

The supernatant was again carefully removed and stored in a 1.5mL Eppendorf tube. Depending on the size of the final pellet, 30-50μL of Diluted Nuclei Buffer (1X Nuclei Buffer, 10X Genomics) was added to the pellet and gently mixed 5 times using a P200 pipette on ice to ensure minimal nuclei clumping. The nuclei suspension was further diluted 1:10 in Diluted Nuclei Buffer prior to quantification. AO/PI staining was performed, and 5μL of the diluted nuclei suspension was loaded into a LUNA-FL™ Dual Fluorescence Cell Counter (Logos Biosystems) for nuclei counting and assesses l lysis completeness and debris.

### Single-nuclei multiomic sequencing (joint snRNA-seq/snATAC-seq)

After counting and quality assessment, nuclei from each sample were prepared for single-cell multiomic sequencing using the protocol outlined in the 10X Genomics Chromium Next GEM Single Cell Multiome ATAC + Gene Expression User Guide Rev A, targeting barcoding of 10,000 nuclei per sample. Transposition Mix from the Chromium Next Gem Single Cell Multiome ATAC + Gene Expression Reagent kit (10X Genomics, #1000283) was added to the nuclei stock, followed by 60 minutes of incubation in a thermal cycler at 37°C. Transposed nuclei were then combined with Master Mix and loaded into the Chromium Next GEM Chip J (10X Genomics, #2000264) along with Single Cell Multiome Gel Beads (10X Genomics, #2000261). The chip was run on the 10X Chromium Controller using the standard Multiome program to 1) generate barcoded, full-length cDNA that is amplified for downstream snRNA-seq gene expression (GEX) library preparation and 2) barcoded transposed DNA for downstream snATAC-seq library preparation.

For both tumor types (siNET and pNET), GEX libraries from all samples were pooled in groups of up to 8 samples and sequenced according to the parameters outlined in the Single Cell Multiome User Guide. snATAC-seq libraries were separately pooled in groups of up to 8 samples and sequenced with appropriate parameters. All single-cell GEX and ATAC-seq libraries were sequenced on S1-100 and SP-100 flowcells using an Illumina NovaSeq 6000 sequencer.

### Spatial transcriptomic sequencing

Hematoxylin and eosin (H&E) slides were prepared from formalin-fixed, paraffin-embedded (FFPE) blocks of all 37 tumors, which were then reviewed by a board-certified pathologist (J.H.) to select high-quality samples for spatial transcriptomic sequencing. Exclusion criteria for FFPE samples consisted of paucity or absence of tumor cells; high density of scar or fibrosis; and tissue degradation, tearing, or shredding.

To determine the quality of RNA recovery from our selected samples, we generated FFPE curls from each tumor block, which were placed in 1.5mL Eppendorf tubes on ice. RNA extraction from each sample’s curls was performed according to the 10x Genomics Visium CytAssist RNA extraction protocol. Agilent Bioanalyzer traces of each sample’s extracted RNA were used to calculate DV200 scores for each sample.

Next, we used the 10x Genomics Visium Spatial Gene Expression protocol to perform spatial transcriptomic sequencing of NET samples whose DV200 scores surpassed 10x Genomics’ recommended thresholds. 10μm sections were cut from each FFPE block and placed on Fisher SuperFrost slides according to 10x Genomics’ Visium CytAssist specifications. The 10x Genomics CytAssist platform was then used to transfer the sectioned tissue to a 6.5 x 6.5mm capture area on 10x Visium Spatial Gene Expression Slides.

Visium Spatial Gene Expression libraries and H&E-stained slides from each sample were generated according to the 10x Genomics protocol; the Gene Expression libraries were then sequenced on an Illumina sequencer.

## Computational Methods

### snRNA-seq and snATAC-seq data preprocessing

Raw base call (BCL) files generated from all single-cell GEX and ATAC-seq libraries were individually demultiplexed and underwent read alignment, barcode counting, peak calling, and cell counting using the standard Cell Ranger ARC analysis pipeline (version 2.0) with default parameters. The pre-built human genome reference included in Cell Ranger ARC (GRCh38-2020-A_arc_v2.0.0) was used for read alignment. Cell Ranger ARC produced raw and filtered count matrices whose rows contained all gene and peak features and whose columns included either all barcodes with non-zero GEX or ATAC-seq signal (raw matrix) or only barcodes identified as cells (filtered matrix).

### snRNA-seq: ambient RNA decontamination, multiplet removal, and cell filtering

Ambient RNA decontamination and empty droplet removal was performed on all samples using the CellBender software package, which uses an unsupervised deep-generative model to parse cell-containing droplets from empty ones to remove technical artifacts from cell-free RNA (39). The raw counts matrix and estimated cell count produced by the Cell Ranger ARC pipeline were inputted into the *remove-background* module of CellBender, and all samples were run using the following default parameters: “ambient” model, low count threshold of 10, learning rate of 0.000001, 150 learning epochs, and false-positive rate of 0.01. For samples whose expected cell counts did not match the CellBender-estimated number of droplets, or whose UMI curve lacked an empty droplet plateau after running CellBender with default parameters, the *--total-droplet-barcodes* parameter was increased to 50,000-75,000. All other samples were run with this parameter set equal to the expected number of cells.

To remove “multiplets,” or technical artifacts caused by unintentional droplet encapsulation of two or more cells, we performed doublet detection and removal on the CellBender-cleaned counts matrix using the Scrublet package (40) (Python, version 0.2.3). Multiplets within the counts matrix were identified and discarded using an expected doublet rate of 0.1-0.2 and manually selected doublet score thresholds. Next, cells with fewer than 200 genes or more than 5% mitochondrial genes were removed from the counts matrix; genes expressed in fewer than 3 cells in the sample were also removed. The resulting ambient RNA-diminished, singlet-only, filtered counts matrix was used for all further downstream quality control and analysis.

### Dimensionality reduction, clustering, and preliminary cell type identification

The count matrix for each sample was log-normalized and scaled using the default *log(x+1)* approach. We then performed linear dimensional reduction on the normalized counts matrix using principal component analysis (PCA) on the top 3,000 highly variable genes across all cells. The first 50 PCA components were used for Leiden clustering (resolution = 0.2). To visualize these preliminary clusters in two-dimensional space, Uniform Manifold Approximation and Projection (UMAP) was performed using the same 50 PCA components. Differential gene expression was performed on a per-cluster basis to identify marker genes and assign preliminary cell type identities for each cluster.

### Across-sample merging, identification of malignant cells, and low-quality cell removal

After preliminary clustering, we concatenated the raw counts matrices and metadata for all 37 samples. This merged object underwent filtering as above to remove cells with <200 counts, genes with non-zero counts in fewer than 3 cells, and cells with > 5% mitochondrial reads. We performed log-normalization and PCA on the top 3,000 highly variable genes, and dimensional reduction as described above. Broad cell type labels were determined by differential gene expression (two-sided Wilcoxon rank-sum test, one cluster vs. cells from all other clusters) on the merged, non-integrated full object.

After we annotate all cells for their corresponding “normal” cell types, we utilized inferCNVpy (version 0.4.5) to identify the malignant cells in each sample (inferCNV of the Trinity CTAT Project). Given the variable proportion of immune cells in each sample, we first generated a uniform “normal cell” reference dataset, by randomly sampling 10% from all immune cells available across the cohort. We used this common reference as the control dataset for all per-sample inferCNVpy runs, and which was run with all mesenchymal cells as the “case” dataset for each sample. inferCNVpy was run on the log-normalized gene expression matrix using a sliding gene window of 250 genes, a CNV score log-fold change clip of 0.1, and a step of 1 (thus computing all possible windows). Then, leiden clustering on the cells’ CNV profiles was conducted on a neighborhood graph computed with 20 neighbors and 20 top PCs at a resolution of 1. Otherwise, default parameters were used. This approach clusters cells based on their inferred copy number alteration profile, and computes a “CNV score” for each of the clusters generated. We defined malignant cells as cells that do not cluster with immune cells (<25% of the cluster are immune cells) and have a CNV score higher than one standard deviation above the mean CNV score of the immune cells. Finally, we performed manual review for each sample to confirm the presence of aberrant copy number alterations in the cells annotated as malignant.

For each sample, non-robustly annotated cells (cNMF annotation != clustering annotation) were removed. A cohort-wide control set was generated by randomly sampling from all immune cells. Neuroendocrine cells were filtered as the case set for each sample. InferCNV was run across all genes (window = 250 genes, log fold change clip = 1, step = 1). InferCNV clusters cells based on their cnv profile and computes cnv score for each cluster. To process InferCNV results, cutoff was defined as mean+1STD cnv score of the control immune cells to filter for cells with significant copy number alterations in each sample’s case set. Results are shown in the CNV classification UMAP for each sample.

### Cell sub-type annotation and visualization

After defining broad cell lineages (e.g., tumor, immune, endothelial), we isolated the raw data corresponding to each cell-type cluster from the integrated object to create new objects for analysis. We then performed *de novo* dimensionality reduction and clustering using the top 3,000 genes and 30 PCs, and Harmony integration was performed on each cell-type object to reduce patient-specific batch effects. To generate cell sub-type annotations, differential gene expression was performed by using a two-sided Wilcoxon rank-sum test to compare expression profiles of one cluster vs. cells from all other clusters.

### Malignant cell program discovery through consensus negative matrix factorization (cNMF) and characterization

To dissect the malignant cell compartment, we employed consensus non-negative matrix factorization (cNMF) (41) per individual patient sample and then aggregated the results as described below. cNMF was performed on a sample-level rather than on the full cohort to avoid detecting patient-specific programs primarily driven by technical factors such as batch effects or copy-number variation (CNV) profiles. For each sample, cNMF was performed on the RNA counts matrix of the 2,000 most highly variable genes, selecting the number of components (k) based on recommended criteria (i.e., inspecting the error and stability plot and picking the smallest k that minimized error while maximizing stability). Density threshold was set to 0.1 for each sample. cNMF programs expressed in too few cells or showing expression of TME-related genes, potentially indicating contamination, were manually removed. The cNMF gene expression programs generated per individual patient sample were characterized by a vector of weights per gene representing its contribution to the program. These programs were combined across all samples by calculating their pairwise cosine similarity after removing small (high score in <10 of cells) or contaminated programs. Hierarchical clustering with an average linkage method was then applied to group similar programs into three clusters for siNETs and four clusters for pNETs. A cNMF program was defined by the median weight of clustered gene expression programs, with the top 200 contributing genes used as a gene signature for the cNMF program. Cells from all patients were scored for the resulting cNMF gene signatures using the scanpy scoring method, i.e., the average gene expression of signature genes subtracted with the average gene expression of control genes. GSEA (42) was run using the prerank function on the ranked list of genes associated with each program, using the hallmarks of cancer as a search database.

### snATAC-seq: quality control and multiplet removal

For each sample, we first filtered the snATAC-seq fragments file produced by CellRanger ARC to contain only barcodes identified by CellRanger as *bona fide* cells. Peak calling was performed using MACS3 (Python, version 3.0.0bl). The resulting peak-by-cell matrix was then used to construct a Signac (43) object (R, version 1.10.0) after filtering for 1) nuclei with a transposition event in at last 200 peaks and 2) peaks detected in at least 10 nuclei.

To identify and remove doublets from the peaks matrix, we scored each nucleus using the scDblFinder method (44) (R, version 1.14.0). Nuclei labeled as “doublet” were removed from the Signac object, and the multiplet-cleaned peaks matrix was carried forward for further processing.

Next, we computed the following five QC metrics for each cell: 1) fraction of fragments in peaks, 2) the ratio of reads in genomic blacklist regions from the ENCODE project, 3) nucleosome signal score, 4) TSS enrichment score, and 5) total number of fragments in peaks. Nuclei with fewer than 1,000 total number of fragments in peaks were removed from the peaks-by-cell matrix. Outlier thresholding for the remaining metrics was determined on a per-sample basis by using the median +/- median absolute deviation (MAD) to set upper and lower bounds.

### snATAC-seq: Dimensionality reduction, clustering, merging, and integration of sample-level datasets

Dimensionality reduction of snATAC-seq data was performed using Signac (43) (R, version 1.11.0). For each sample, term frequency-inverse document frequency (TF-IDF) normalization was performed on all cells passing QC thresholds. All features were used to run singular value decomposition (SVD) on the normalized matrix. LSI dimensions 2 through 30 were used for UMAP projection and downstream clustering; the first LSI dimension correlated strongly with sequencing depth and was not included. Shared nearest neighbor (SNN)-based clustering and differential gene activity were performed to identify broad cell types within each sample. For samples containing more than one cell type, peaks were called on a cluster-level basis using MACS3; for samples containing only one cluster, bulk peaks were called using MACS3.

To merge snATAC-seq data from the sample-level datasets, we first created a common set of peaks using the GenomicRanges *reduce* function on the cluster-level or bulk-level peaks from each sample. These peaks were quantified and used to create new Seurat objects for each dataset. Next, we merged all Seurat objects per the default Signac workflow. Gene activity was inferred for all nuclei in the merged dataset using the *GeneActivity* function from Signac. Cell type annotations from matched snRNA-seq nuclei and differential gene activity between snATAC-seq clusters were combined to determine broad cell type annotations.

To refine peak representation within the merged object, broad cell types were used to re-call peaks on the integrated object with MACS3. These refined peaks were then used to construct a new Signac object, and low-quality cells were again excluded using the MAD-based thresholds above (peak region fragments > 1000). Broad cell types were again annotated using a combination of differential gene activity and cell type labels from matched snRNA-seq metadata. The final Signac object contained 74,004 nuclei from 35 samples (20 PNETs, 15 SINETs).

### Gene regulatory network analysis and visualization

To analyze gene regulatory networks (GRNs) in malignant pNET cells, we utilized the Python-based single-cell regulatory network inference and clustering (pySCENIC) pipeline (45). To minimize confounding by unequal distribution of cell types, we first created 10 downsampled objects from the snRNA-seq data containing a maximum of 200 cells per Symphony-mapped cell type (i.e. alpha, beta, gamma, delta, or epsilon). From each downsampled object, we generated a log-normalized counts matrix containing all tumor cells containing <5% mitochondrial reads and greater than 200 genes, and all genes with non-negative counts in at least 3 cells. Mitochondrial and long-noncoding RNA genes were removed from the counts matrix prior to normalization to avoid quantification of regulons associated with those genes.

GRNBoost2 (*Arboreto* multiprocessing implementation) was then used to perform GRN inference on each of the 10 downsampled objects. For each object, the resulting adjacencies matrix was inputted into pySCENIC’s cisTarget pipeline for regulon prediction (motif databases: *hg38_10kbp_up10kbp_down* and *hg38_500bp_up_100bp_down*; ranking database: *motifs-v10nr_clust-nr.hgnc-m0.001-o0.0*). Regulons that were identified in at least 8 of the 10 pySCENIC runs were considered robust; all other regulons were discarded. Finally, AUCell was used to calculate the enrichment of each robust regulon per individual cell in the full pNET malignant cell dataset. To visualize the top-ranked regulons per relevant grouping (e.g., sample, site, etc.), we first generated z-scores from each regulon’s AUC scores. We then selected the top 5 regulons per subgroup and plotted their standardized scores using Seaborn’s *clustermap* function.

### Motif enrichment analysis

Transcription factor (TF) motif enrichment analysis of snATAC-seq data was performed using *chromVAR* per the Signac documentation (43,46). Briefly, cell type labels were assigned to cells using matched barcodes from snRNA-seq analysis. Motif DNA sequence information from the JASPAR2020 collection was added to the Signac object using the hg38 genome (47). Motif activity per cell was calculated by running the *RunChromVAR* function (Signac) with default parameters and the hg38 genome; enriched motifs per broad cell type were identified using the *FindAllMarkers* function (Signac).

### Spatial transcriptomics sequencing data pre-processing

Processing of Illumina sequencing output of all CytAssist Spatial Gene Expression for FFPE libraries was performed using the 10x Genomics Space Ranger workflow. Raw Illumina BCL files were demultiplexed and converted to FASTQ files using the *spaceranger mkfastq* pipeline (version 2.1.0). Next, the *spaceranger count* pipeline was used to perform fiducial image alignment, tissue detection, read alignment (Space Ranger reference genome: GRCh38-2020-A), and barcode and UMI counting. Manual registration of the CytAssist and H&E image files was performed using the 10x Genomics Loupe platform. The filtered counts matrix produced by Space Ranger was used for downstream quality control and analysis.

### Spatial transcriptomics quality control, dimensionality reduction, and clustering

Analysis of spatial transcriptomics data was performed using Squidpy (48) (Python, version 1.3.1). Visium spots with fewer than 5,000 or greater than 35,000 counts or with greater than 20% mitochondrial reads were filtered out of the Space Ranger-filtered counts matrix. The remaining spots were normalized using SCTransformPy (https://github.com/atarashansky/SCTransformPy), and the top 5,000 highly variable genes were used for downstream PCA, UMAP, and Leiden clustering analysis (number of PCs = 50). cNMF programs were scored using AUCell scoring on the cancer cell compartments of each of the spatial transcriptomics samples.

To assess the spatial localization of malignant programs, we calculated the distance of each tumor spot to the tumor boundary in the Visium data. Malignant spots were classified as cNMF1-high or cNMF2-high using sample-specific program score thresholds, particularly taking the top 10th percentile for each. Euclidean distances from each spot to the nearest tumor–normal boundary were computed from Visium spatial coordinates, generating a dist_to_boundary measure for every spot. To account for differences in tumor size across samples, distances were normalized by the maximum distance within each sample, yielding a core-periphery index (0 = tumor edge, 1 = deep core).

To quantify how malignant programs spatially associate with specific microenvironmental compartments, we computed the distance from each cNMF1-high or cNMF2-high tumor spot to the nearest immune, endothelial, or fibroblast spot in the Visium arrays. Distances were converted into “ring units,” defined as the mean nearest-neighbor spacing within each sample. Spots were then classified as near a given cell type if they fell within preset ring thresholds (≤1 ring). For each sample and malignant program, we calculated the fraction of tumor spots falling within these proximity thresholds. Differences between cNMF1-high and cNMF2-high tumor spots were evaluated using paired Wilcoxon signed-rank tests applied to per-sample fractions. Proximity fractions were visualized using barplots comparing program-specific associations across samples.

### Cell-cell communication analysis

Cell-cell communication was inferred using LIANA (v1.5.1) with the CellChat statistical framework and CellChatDB (29,49). Normalized single-nucleus RNA-seq data were grouped by pathology subtype, and ligand-receptor interactions were scored for communication probability and statistical significance. Significant interactions were visualized with LIANA’s dot plot to identify program-specific signaling patterns.

### Link between malignant programs and clinical characteristics

To explore whether the identified cNMF programs correlated with clinical characteristics in our cohort, we assessed the relationship between program scores and various clinical characteristics. We subsetted the cancer spots from our single cell data and performed program scoring for both siNET and pNET. We then normalized these scores to the number of patients in each group.

### Link between malignant programs and clinical characteristics in external bulk datasets

To validate our findings, we assessed the scores of these programs in external bulk datasets and examined their association with various clinical parameters. RNA-seq expression data and clinical characteristics from the study by Greenburg et al. were downloaded, with the RNA-seq raw counts transformed into transcript per million (TPM) (37). The cNMF programs were scored using single-sample Gene Set Enrichment Analysis (ssGSEA) (42). The resulting scores were then correlated with TNM staging and metastatic status.

### Statistics

Statistical analyses were performed using R and Python. All statistical tests were two-sided, and a P value < 0.05 was considered statistically significant unless otherwise indicated. Sample sizes were determined by the availability of high-quality patient specimens. Comparisons between groups were performed using non-parametric tests unless otherwise noted, and multiple hypothesis testing was controlled using the Benjamini–Hochberg false discovery rate procedure. Additional details are provided in the figure legends.

## AUTHOR CONTRIBUTIONS

J.K.: conceptualization, investigation, software, formal analysis, visualization, writing – original draft; S.E.H: conceptualization, data curation, investigation, software, formal analysis, writing – original draft, writing – review & editing; A.G.: investigation, writing – review & editing; D.G.: investigation, writing – review & editing; H.I.H.: investigation, writing – review & editing; B.M.T: investigation, writing – review & editing; Y.T.: investigation, writing – review & editing; E.P.: investigation, writing – review & editing; T.P.: investigation, writing – review & editing; L.V.: investigation; K.B.: investigation; R.G.: investigation; L.B.: resources; E.S.: project administration; J.L.H.: resources; J.P.: conceptualization, investigation, writing – review & editing; J.C.: conceptualization, funding acquisition, resources, data curation, supervision, writing – review & editing; E.M.V.A: conceptualization, funding acquisition, resources, supervision, writing – original draft, writing – review & editing

## FUNDING

This work was supported by:

The Orr Family Foundation (J.K., S.E.H., J.C., E.M.V.A.)

NIH 5U2CCA233195-05 (E.M.V.A.)

NIH R37 CA222574 (J.P., E.M.V.A.)

NIH R01 CA227388 (J.P., E.M.V.A.)

The Goldhirsh-Yellin Foundation (J.C.)

The project described was supported by award Number T32GM144273 (S.E.H.) from the National Institute of General Medical Sciences. The content is solely the responsibility of the authors and does not necessarily represent the official views of the National Institute of General Medical Sciences or the National Institutes of Health.

## ACKNOWLEDGEMENTS

We thank the patients who participated in this study. The authors would also like to thank the following core facilities: the Single Cell Core at Harvard Medical School (Boston, MA) for performing the single-nuclei RNA-/ATAC-seq library generation; the Molecular Biology Core Facilities at Dana-Farber Cancer Institute (Boston, MA) for performing the Illumina sequencing of snRNA/ATAC-seq libraries; the Broad Institute Genomics Platform (Cambridge, MA) for performing the sample preparation and sequencing for whole-exome and bulk RNA-sequencing; and the Specialized Histopathology Core at Brigham and Women’s Hospital (Boston, MA) and the Spatial Technologies Unit (Boston, MA) for spatial transcriptomics sample preparation and sequencing.

