## Supplemental Figures for "Conserved Neuronal-like and Secretory Programs Define the Spatial Architecture of Gastroenteropancreatic Neuroendocrine Tumors"

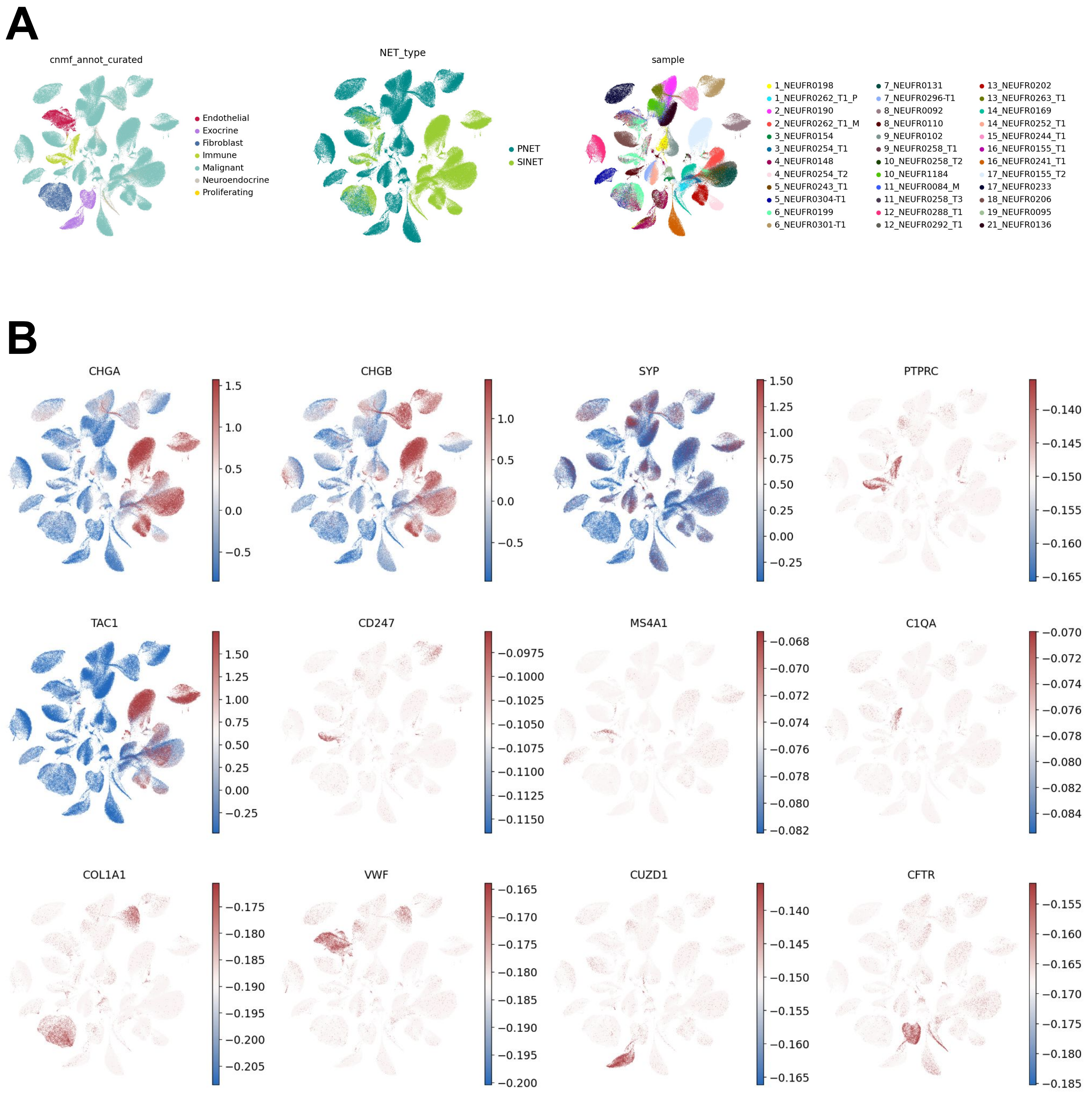


**Supplementary Figure 1: Cell type annotation and malignant cell identification in single-nucleus multi-omic data.** (A) UMAP embedding of snRNA-seq data colored by broad cell type annotations derived from Leiden clustering and canonical marker gene expression, including malignant cells, immune populations, endothelial cells, cancer-associated fibroblasts (CAFs), and normal epithelial and exocrine cells. (B) Identification of malignant cells using inferred copy number alteration (CNA) profiles combined with expression of established neuroendocrine tumor marker genes. Heatmaps display large-scale chromosomal gains and losses across malignant and non-malignant populations. CNA inference was performed as described in Methods and used to distinguish malignant tumor cells from surrounding stromal and immune cells.

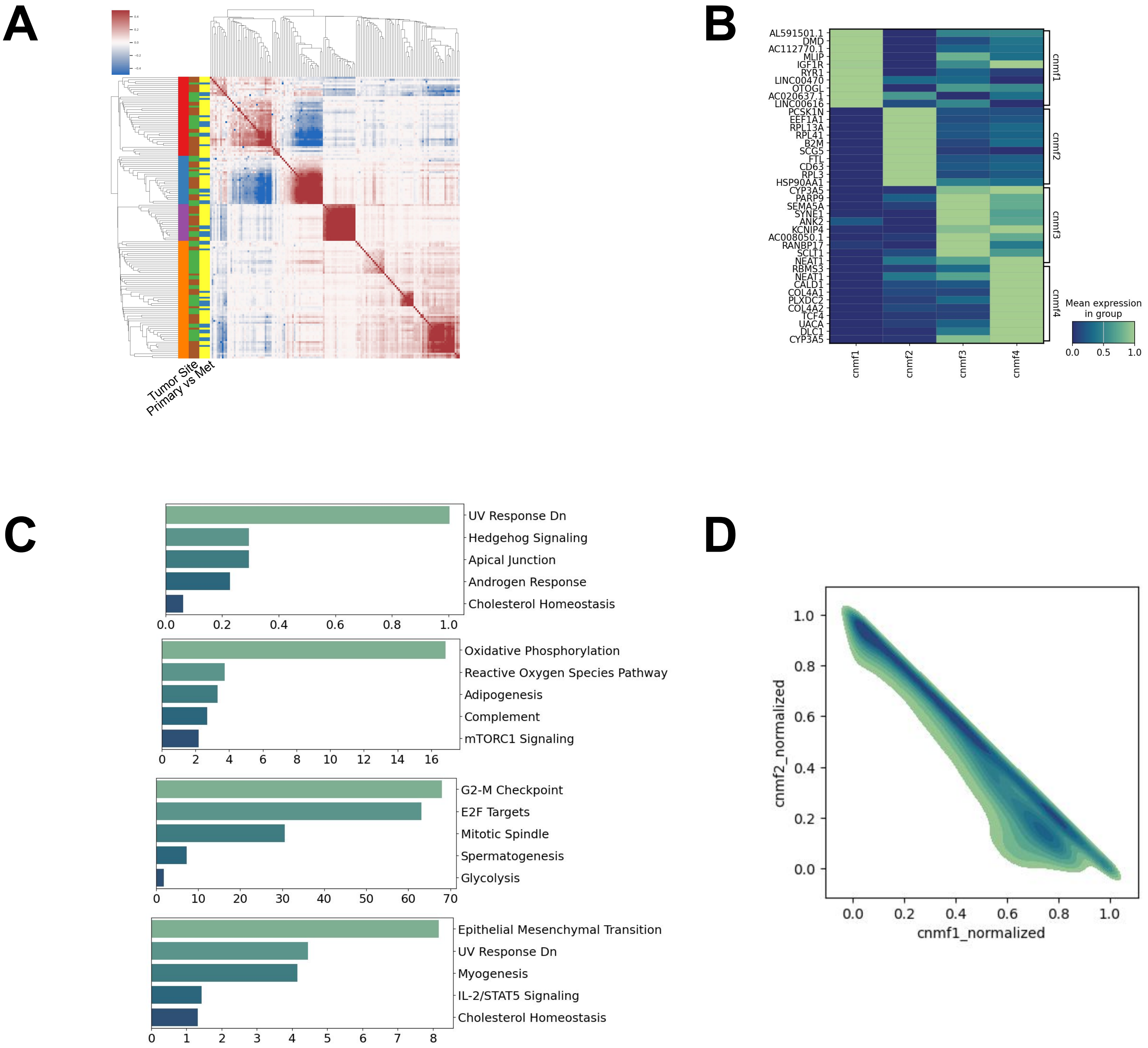


**Supplementary Figure 2: Integrated cNMF analysis reveals conserved malignant transcriptional programs across tumor types.** Integrated consensus non-negative matrix factorization (cNMF) was performed on pooled malignant cells from siNETs and pNETs. (A) Heatmap of gene weights for integrated cNMF programs, demonstrating overlap with programs identified independently in siNETs and pNETs. The programs are also independent of primary vs metastatic samples. (B) Contribution of siNET- and pNET-derived malignant cells to each integrated cNMF program, illustrating that both tumor types contribute to neuronal-like and secretory neuroendocrine programs. (C) Density plots of normalized integrated cNMF program scores showing continuous distributions rather than discrete subpopulations. These analyses support the presence of conserved malignant cell states across organ sites.

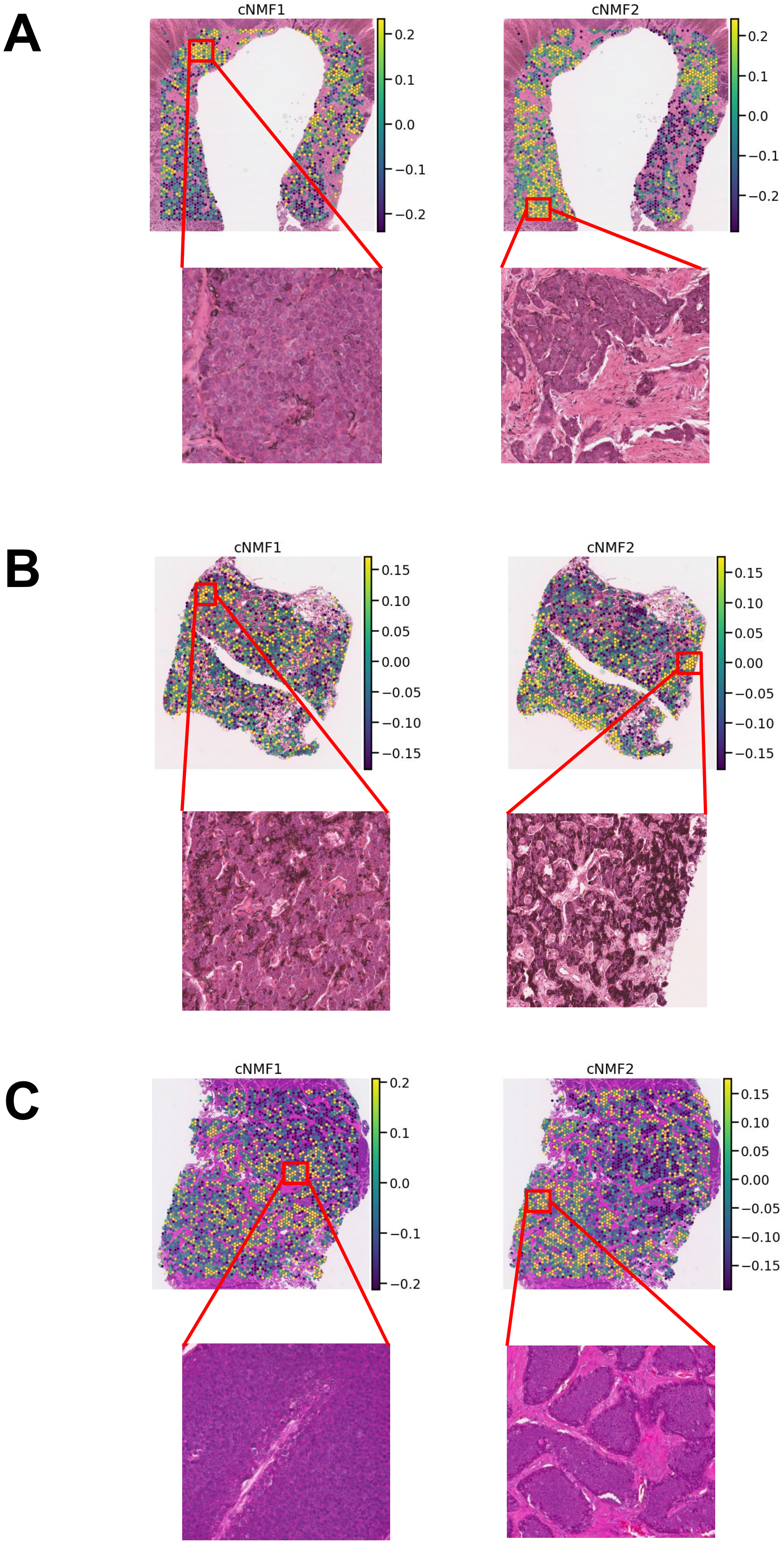
A
cNMF1
0.2
0.1
0.0
−0.1
−0.2
cNMF2
0.2
0.1
0.0
−0.1
−0.2
B
cNMF1
0.15
0.10
0.05
0.00
−0.05
−0.10
−0.15
cNMF2
0.15
0.10
0.05
0.00
−0.05
−0.10
−0.15
C
cNMF1
0.2
0.1
0.0
−0.1
−0.2
cNMF2
0.15
0.10
0.05
0.00
−0.05
−0.10
−0.15

**Supplementary Figure 3: Additional spatial examples validating localization of malignant cNMF programs.** cNMF1 and cNMF2 scoring of malignant spots on spatial transcriptomics data with representative spatial transcriptomic sections demonstrating localization of cNMF1-high malignant spots to densely cellular tumor regions and cNMF2-high malignant spots to stromal-infiltrated regions. (A) Sample S13, (B) Sample S8-T2, (C) S4. Across samples, spatial enrichment patterns were consistent with those shown in Figure 4. These panels provide qualitative validation across multiple tumors; statistical analyses of spatial proximity and boundary distance are presented in the main figures.

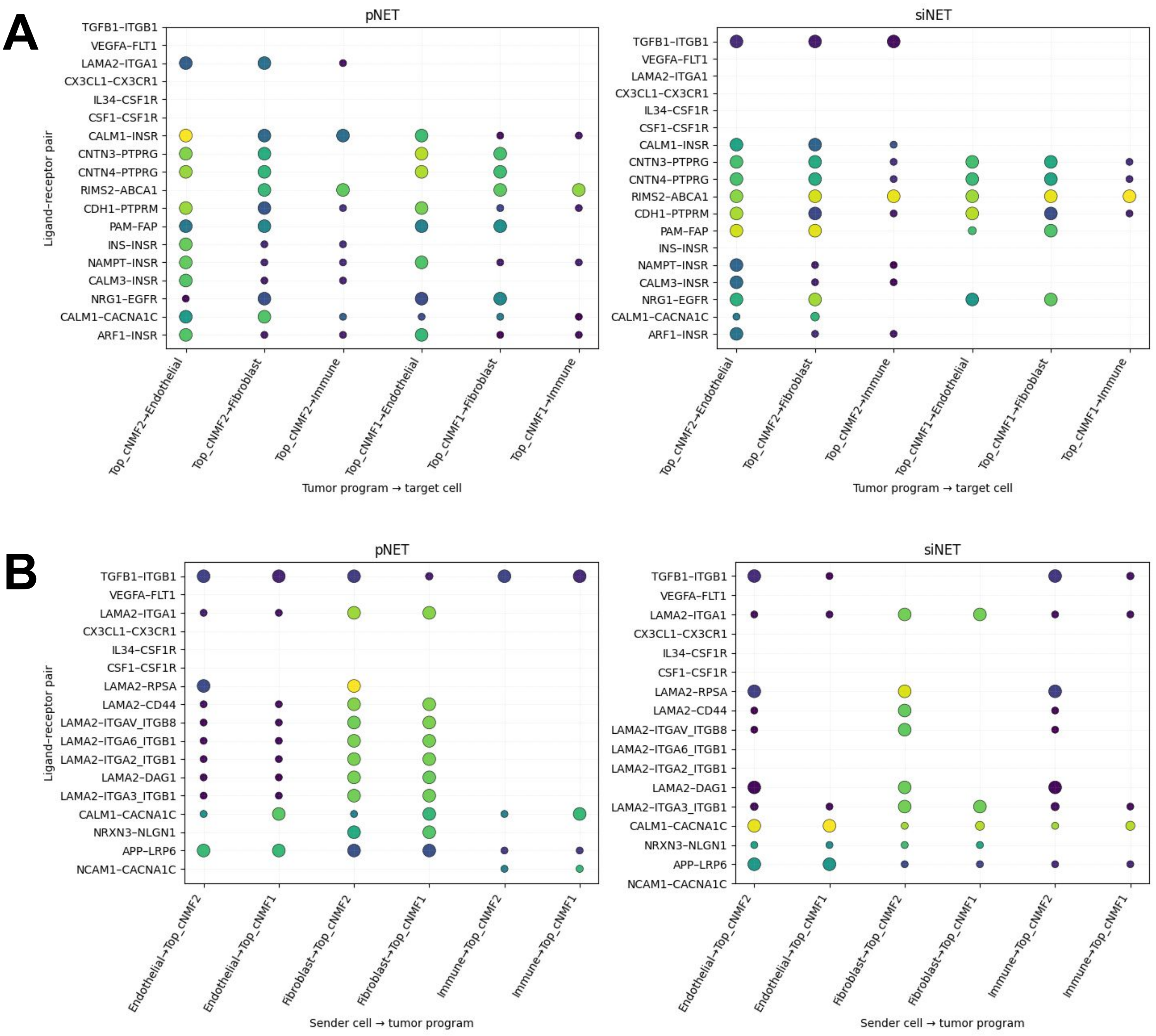


**Supplementary Figure 4. Bidirectional ligand–receptor signaling associated with malignant cell programs.** Bubble plots show prioritized ligand–receptor interactions involving Top_cNMF1 and Top_cNMF2 malignant cell programs in pancreatic and small intestinal neuroendocrine tumors (pNETs and siNETs). (A) Outgoing signaling from malignant programs to stromal and immune populations reveals enriched endothelial and fibroblast interactions from Top_cNMF2 malignant cells, whereas Top_cNMF1 malignant cells preferentially engage immune and myeloid populations. (B) Incoming signaling from stromal and immune populations to malignant programs demonstrates reciprocal patterns, with Top_cNMF2 malignant cells receiving prominent vascular and stromal signals and Top_cNMF1 malignant cells receiving immune- and neuronal-associated signals. Dot size represents interaction significance (−log10 p-value), and dot color represents interaction strength as quantified by LIANA.

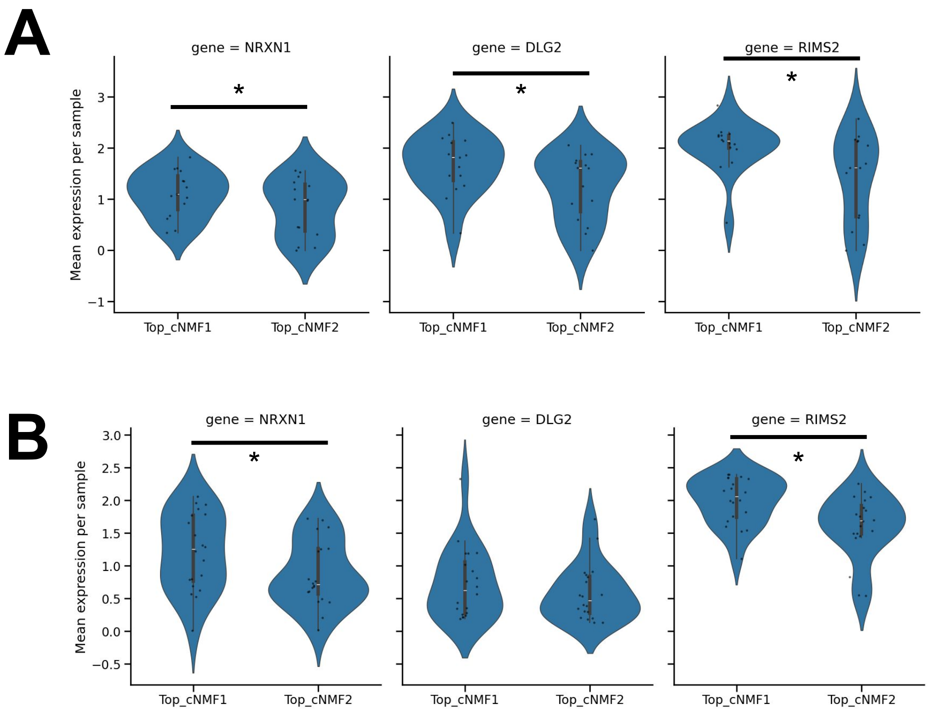


**Supplementary Figure 5: Neuronal communication–associated genes are enriched in si/p-cNMF1-high malignant cells.** Violin plots show per-sample mean expression of the neuronal communication–associated genes NRXN1, DLG2, and RIMS2 in malignant cells stratified by Top-cNMF program (Top-cNMF1 vs. Top-cNMF2). (A) siNET samples were scored for si-cNMF1 amd si-cNMF2. (B) pNET samples were scored for p-cNMF1 and p-cNMF2. For each sample, gene expression was averaged across all malignant cells assigned to the indicated Top-cNMF program. Black points represent individual samples. Statistical significance was assessed using a paired two-sided Wilcoxon signed-rank test across samples, with p-values adjusted for multiple testing using the Benjamini–Hochberg method. * = p < 0.001

A

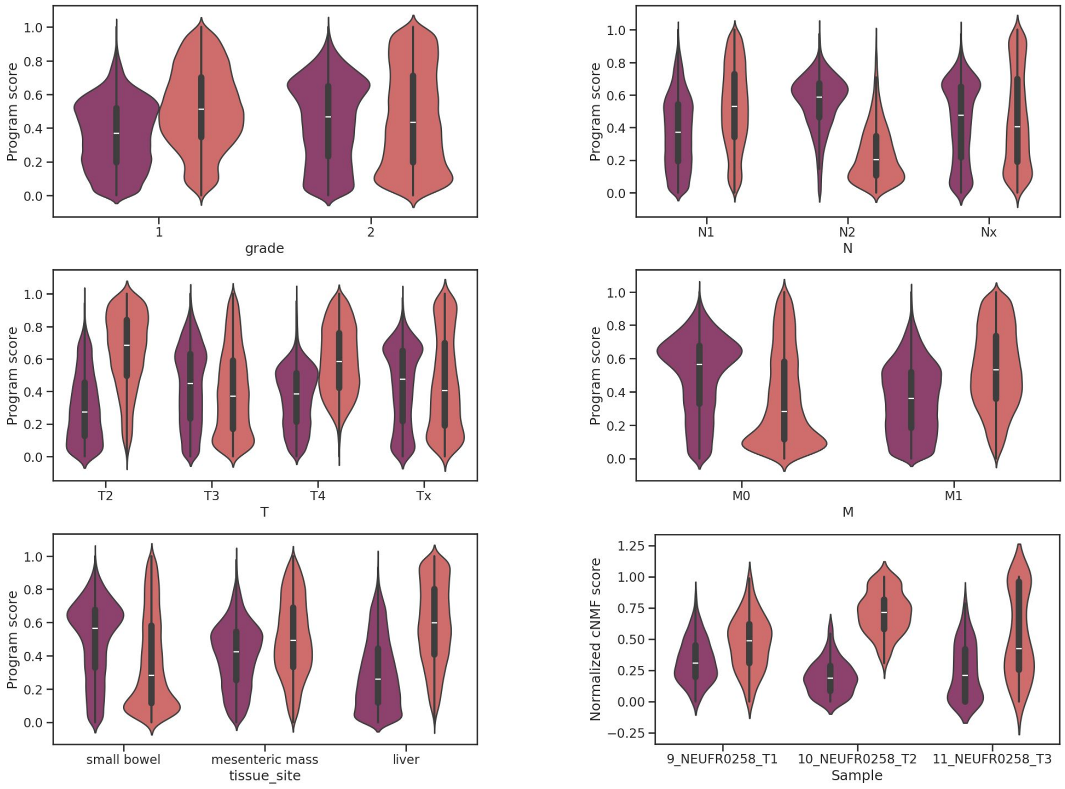
Program score
grade
1
2
N1
N2
Nx
N
T2
T3
T4
Tx
T
M0
M1
M
small bowel
mesenteric mass
liver
tissue_site
Normalized cNMF score
9_NEUFR0258_T1
10_NEUFR0258_T2
11_NEUFR0258_T3
Sample

B

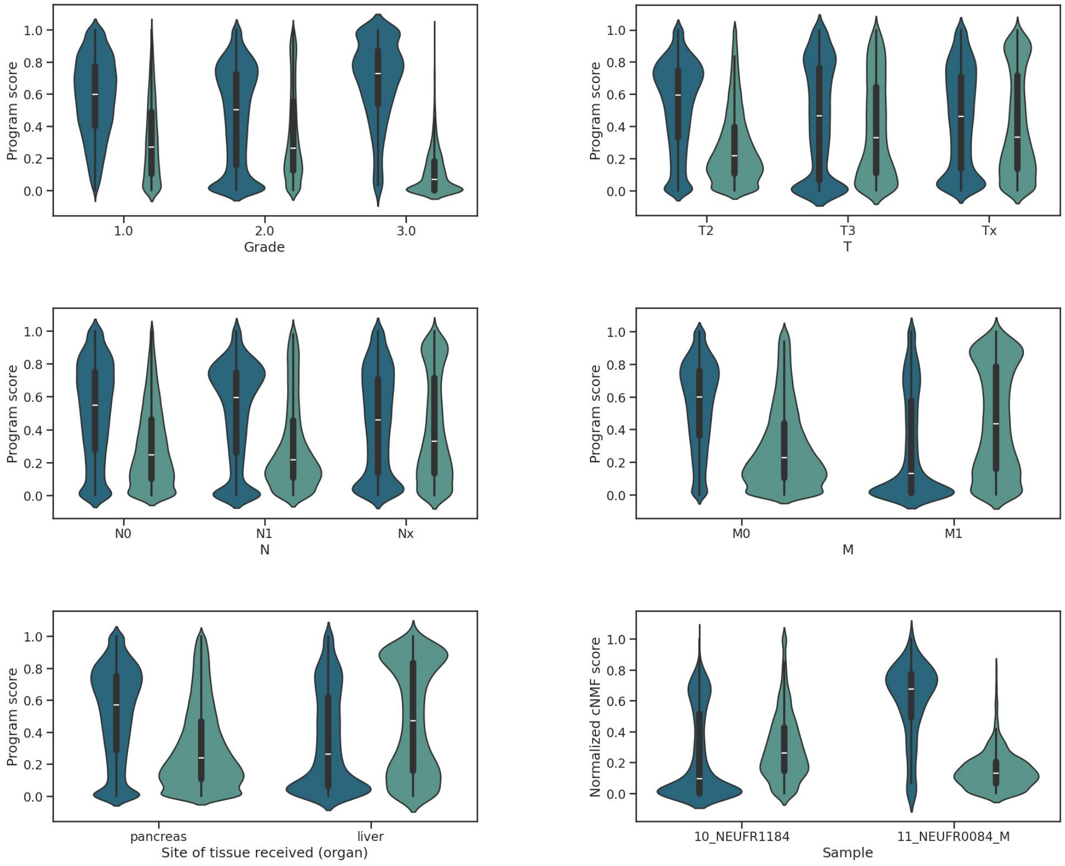
Program score
1.0
2.0
3.0
Grade
T2
T3
Tx
T
N0
N1
Nx
N
M0
M1
M
pancreas
liver
Site of tissue received (organ)
Normalized cNMF score
10_NEUFR1184
11_NEUFR0084_M
Sample

**Supplementary Figure 6. Malignant cNMF program scores associate with metastatic status and progression.**
(A) Distribution of si-cNMF program scores stratified by Grade, N, T, M, and tissue site across siNETs. Within-patient analysis of a representative siNET case (sample S8) sampled across primary, regional metastatic, and distant metastatic lesions. Statistical significance was assessed using a two-sided Wilcoxon rank-sum test across samples. (B) Distribution of p-cNMF program scores stratified by Grade, N, T, M, and tissue site across pNETs. Statistical significance was assessed using a two-sided Wilcoxon rank-sum test across samples.
